# Identification of the myogenetic oligodeoxynucleotides (myoDNs) that promote differentiation of skeletal muscle myoblasts by targeting nucleolin

**DOI:** 10.1101/2020.10.07.330472

**Authors:** Sayaka Shinji, Koji Umezawa, Yuma Nihashi, Shunichi Nakamura, Takeshi Shimosato, Tomohide Takaya

## Abstract

Herein we report that the 18-base telomeric oligodeoxynucleotides (ODNs) designed from the *Lactobacillus rhamnosus* GG genome promote differentiation of skeletal muscle myoblasts which are myogenic precursor cells. We termed these myogenetic ODNs (myoDNs). The activity of one of the myoDNs, iSN04, was independent of Toll-like receptors, but dependent on its conformational state. Molecular simulation and iSN04 mutants revealed stacking of the 13-15th guanines as a core structure for iSN04. The alkaloid berberine bound to the guanine stack and enhanced iSN04 activity, probably by stabilizing and optimizing iSN04 conformation. We further identified nucleolin as an iSN04-binding protein. Results showed that iSN04 antagonizes nucleolin, increases the levels of p53 protein translationally suppressed by nucleolin, and eventually induces myotube formation by modulating the expression of genes involved in myogenic differentiation and cell cycle arrest. This study shows that bacterial-derived myoDNs serve as aptamers and are potential nucleic acid drugs directly targeting myoblasts.

## Introduction

Skeletal muscle myoblasts are myogenic precursor cells that play a central role during muscle development and regeneration. In the first step of these processes, muscle stem cells called satellite cells on myofibers are activated into myoblasts. After several rounds of division, myoblasts differentiate into myocytes, which is led by myogenic transcription factors such as MyoD and myogenin. Myocytes then fuse to form multi-nuclear myotubes to generate or restore myofibers (Dumont et al., 2015). However, the differentiation ability of myoblasts declines due to aging or diseases. Aged murine myoblasts tend to differentiate into the fibrogenic lineage by the activation of the canonical Wnt pathway (Brack et al., 2007). The myoblasts isolated from the chronic kidney disease mice model showing muscle atrophy display attenuated MyoD expression and myotube formation (Zhang et al., 2010). Cancer-conditioned media inhibit myogenic differentiation by upregulating C/EBPβ in the murine myoblast cell line C2C12 (Marchildon et al., 2015). Such hypoactivities of myoblasts are considered to be one of the reasons for the development of muscle atrophy (Fukada, 2018; McCormick and Vasilaki, 2018). Therefore, myoblast differentiation can be a clinical target for sarcopenia, disease-related muscle wasting, and cancer cachexia which are risk factors for mortality (Anker et al., 1997; Rubin, 2003; Carrero et al., 2008).

Several molecules have been identified that facilitate myogenic differentiation. Histone deacetylase inhibitors (HDACIs), trichostatin A (TSA) and valproic acid (VPA), promote myotube formation by inducing follistatin in myoblasts (Iezzi et al., 2004). However, TSA and VPA are non-specific HDACIs that affect a broad range of biological processes in vivo. Recent studies have reported that the combined treatment of ursolic acid (UA) with leucine (Kim et al., 2015), and a single treatment of an oleanolic acid (OA) derivative (Cui et al., 2019) potentiates differentiation of C2C12 cells. As the half-lives of UA in plasma and OA in serum are less than 1 h (Chen et al., 2011; Li et al., 2012), their pharmacokinetic parameters need to be improved for clinical application.

Nucleic acids have tremendous potential for use in next-generation drugs. They are chemically synthesized, stable, and modifiable molecules that can access diverse targets with high specificities. Complementary antisense oligonucleotides modulate gene expression by degrading mRNAs, trapping microRNAs, or correcting splicing events (Quemener et al., 2020). Other types of oligonucleotides serve as aptamers that specifically interact with their target proteins (Wang et al., 2019). Furthermore, many immunomodulatory oligodeoxynucleotides (ODNs) from microbial and autologous DNA sequences have been reported. ODNs with unmethylated CpG motifs (CpG-ODNs) serve as ligands for Toll-like receptor (TLR) 9 and initiate an inflammatory cascade (Vollmer and Krieg, 2009). In contrast, inhibitory ODNs (iODNs) representatively expressing telomeric elements suppress immunological reactions depending on TLR3, TLR7, and TLR9 (Klinman et al., 2008; Sackesen et al., 2013). At present, CpG-ODNs and iODNs are anticipated to be effective drugs for sepsis and allergic diseases (Yamamoto et al., 2017; Wang et al., 2015).

Intriguingly, some CpG-ODNs have been reported to alter cell fate. Initial studies have shown that CpG-1826 modulates osteoclastogenesis through TLR9 (Zou et al., 2003; Amcheslavsky et al., 2005). CpG-KSK13 displayed an inhibitory effect on osteoclast differentiation by downregulating TREM-2 (Chang et al., 2009). CpG-2006 and its variants interfere with osteoblast differentiation from mesenchymal stem cells (MSCs) by inhibiting the BMP-Smad signal in a TLR9-independent manner (Norgaard et al., 2010). By contrast, MT01, a 27-base C-rich iODN (Yang et al., 2010), stimulates the differentiation of MSCs into osteoblasts via the ERK-p38 pathway (Feng et al., 2011; Shen et al., 2012; Hou et al., 2012).

These findings prompted us to explore a novel ODN that regulates myoblast differentiation. We recently constructed 18-base ODN candidates designed from the genome sequence of a lactic acid bacteria strain, *Lactobacillus rhamnosus* GG (LGG) (Nigar et al., 2017). These synthetic phosphorothioated (PS)-ODNs resistant to nucleases were applied to myoblasts to validate their myogenetic effects. Herein, we report a series of 18-base telomeric PS-ODNs, named myogenetic ODNs (myoDNs), that promote myoblast differentiation depending on their conformation but independent of TLR signal. This study presents an innovative approach to regulate cell fate using bacterial-derived ODNs.

## Materials and Methods

### ODNs and Chemicals

The sequences of the ODNs used in this study are described in Supplementary Table S1. PS-ODNs, 6-carboxyfluorescein (6-FAM)-conjugated PS-ODNs, and biotin-conjugated PS-ODNs were synthesized and purified via HPLC (GeneDesign, Osaka, Japan). AS1411 having a phosphodiester backbone was synthesized and desalted (Integrated DNA Technologies, Coralville, IA, USA) as previously reported (Girvan et al., 2006). PS-ODNs, AS1411, berberine hydrochloride (Nacalai, Osaka, Japan), palmatine chloride hydrate (Nacalai), coptisine chloride (Wako, Osaka, Japan), and jatrorrhizine (Wako) were dissolved in endotoxin-free water. An equal volume of endotoxin-free water instead of PS-ODNs and berberine analogs served as negative controls.

### Cell Culture

All cells were cultured at 37°C under 5% CO2 throughout the experiments.

Murine myoblasts (mMBs) were isolated from the skeletal muscle of 4-week-old C57BL/6J mice (Clea Japan, Tokyo, Japan) and primary-cultured as previously described (Takaya et al., 2017; Nihashi et al., 2019b). mMBs were maintained on the dishes or plates coated with collagen type I-C (Cellmatrix; Nitta Gelatin, Osaka, Japan), and cultured in growth medium (GM) for mMB consisting of Ham’s F10 medium (Thermo Fisher Scientific, MA, USA), 20% fetal bovine serum (FBS) (HyClone; GE Healthcare, UT, USA), 2 ng/ml recombinant human basic fibroblast growth factor (Wako), and a mixture of 100 units/ml penicillin and 100 μg/ml streptomycin (PS) (Nacalai).

Primary-cultured human myoblast (hMB) stock of adult healthy female (CC-2580; Lonza, MD, USA) was maintained according to the manufacturer’s instruction. hMBs were seeded on collagen-coated dishes, cultured in Skeletal Muscle Growth Media-2 (CC-3245; Lonza) as GM for hMB, and differentiation was induced in differentiation medium (DM) for hMB consisting of DMEM (Nacalai) with 2% horse serum (HS) (HyClone; GE Healthcare) and PS.

C2C12 cells (DS Pharma Biomedical, Osaka, Japan) were seeded on collagen-coated dishes, cultured in GM for C2C12 cells consisting of DMEM with 10% FBS and PS, and induced differentiation in DM for C2C12 cells consisting of DMEM with 2% HS and PS.

Murine embryonic fibroblasts (MEFs) were prepared from E12 embryos of Slc:ICR mice (Japan SLC, Shizuoka, Japan). The embryos that were removed their heads and internal organs were minced in GM for MEFs consisting of DMEM with 10% FBS and PS. The tissue clusters were seeded and cultured for 3 days. Then outgrown cells were dissociated to single cells as MEFs using 0.25% trypsin with 1 mM EDTA (Wako).

MC3T3-E1 cells (RCB1126) were provided by RIKEN BRC (Tsukuba, Japan) through the Project for Realization of Regenerative Medicine and the National Bio-Resource Project of the MEXT, Japan. The cells were maintained in EMEM (Wako) with 10% FBS and PS.

### Immunocytochemistry

Immunocytochemistry of myoblasts was performed as previously described (Takaya et al., 2017; Nihashi et al., 2019a; Nihashi et al., 2019b). The myoblasts were fixed with 2% paraformaldehyde, permeabilized with 0.2% Triton X-100, and immunostained with 0.5 μg/ml mouse monoclonal anti-myosin heavy chain (MHC) antibody (MF20; R&D Systems, MN, USA) and 1.0 μg/ml rabbit polyclonal anti-nucleolin antibody (ab22758; Abcam, Cambridge, UK). 0.1 μg/ml each of Alexa Fluor 488-conjugated donkey polyclonal anti-mouse IgG antibody and Alexa Fluor 594-conjugated donkey polyclonal anti-rabbit IgG antibody (Jackson ImmunoResearch, PA, USA) were used as secondary antibodies. Cell nuclei were stained with DAPI (Nacalai). High-resolution fluorescent images were taken under an EVOS FL Auto microscope (AMAFD1000; Thermo Fisher Scientific). The ratio of MHC^+^ cells was defined as the number of nuclei in the MHC^+^ cells divided by the total number of nuclei, and the fusion index was defined as the number of nuclei in the multinuclear MHC^+^ myotubes divided by the total number of nuclei using ImageJ software (National Institutes of Health, USA).

### Screening System

1.0×10^4^ mMBs or 5.0×10^3^ hMBs in 100 μl GM/well were seeded on collagen-coated 96-well plates. The next day, the medium was replaced with GM for mMB or DM for hMB containing PS-ODNs. After 48 h, the mMBs or hMBs were subjected to MHC and DAPI staining. Fluorescent images were automatically captured using CellInsight NXT (Thermo Fisher Scientific). The ratio of MHC^+^ cells of mMBs and MHC signal intensities of hMBs were automatically measured using HCS Studio: Cellomics Scan software (Thermo Fisher Scientific). The average value of three wells (4 fields/well) served as the mean of each sample.

### Cell Counting

5.0×10^4^ mMBs/well were seeded on collagen-coated 24-well plates and 5.0×10^4^ MEFs/well were seeded on 12-well plates. The next day, the medium was replaced with medium containing 1 or 3 μM iSN04. The cells were continuously cultured until cell counting. For counting, the cells were completely dissociated using 0.25% trypsin with 1 mM EDTA and the number of cells was counted using a hemocytometer.

### Quantitative real-time RTPCR (qPCR)

2.5×10^5^ hMBs in GM were seeded on collagen-coated 60-mm dishes. The next day, the medium was replaced with DM containing 30 μM iSN04. After 24 h, total RNA of the hMBs was isolated using NucleoSpin RNA Plus (Macherey-Nagel, Düren, Germany) and was reverse transcribed using ReverTra Ace qPCR RT Master Mix (TOYOBO, Osaka, Japan). qPCR was performed using GoTaq qPCR Master Mix (Promega, WI, USA) with StepOne Real-Time PCR System (Thermo Fisher Scientific). The amount of each transcript was normalized to that of tyrosine 3-monooxygenase/tryptophan 5-monooxygenase activation protein zeta gene (*YWHAZ*). The results are presented as fold-change. Primer sequences are listed in Supplementary Table S2.

### RNA sequencing (RNAseq)

The total RNA of hMBs used for qPCR was subjected to RNA-seq (Novogene, Beijing, China). RNA quality was checked using an Agilent 2100 Bioanalyzer (Agilent Technologies, Waldbronn, Germany). RNA integrity number (RIN) values were 10.0 (max score) in all samples (Supplementary Figure S4A). The RNA was subjected to library preparation using Illumina TruSeq RNA and DNA Sample Prep Kits (Illumina, CA, USA). Library quality was confirmed using a Qubit 2.0 fluorometer (Life Technologies; Thermo Fisher Scientific) and Agilent 2100 Bioanalyzer. RNA-seq was performed using Illumina NovaSeq 6000 (Illumina) to generate > 6-GB raw data per sample. Raw data were recorded in FASTQ format. The quality of the read was calculated as the arithmetic mean of the Phred quality score. The reads with following characteristics were discarded: adapter contamination, when uncertain nucleotides constituted > 10% of either read, or when low quality nucleotides (base quality < 20) constituted > 50% of the read. The cleaned reads were mapped to a human reference genome (GRCh38.82) using TopHat2. The number of the reads and mapping efficiencies are summarized in Supplementary Table S3. Expression levels of the transcripts were calculated as fragments per kilobase per million reads (FPKM) values using HTSeq. False discovery rate (FDR) was employed to correct their *p* values.

### Heatmap

Heatmaps of FPKM values were generated via Heatmapper (http://www.heatmapper.ca/) (Babicki et al., 2016) with the following settings: Clustering method, Average linkage; Distance measurement method, Pearson.

### Gene Ontology (GO)Analysis

The iSN04-dependent differentially expressed genes (DEGs) were subjected to GO analysis using DAVID Bioinformatics Resources 6.8 (https://david.ncifcrf.gov/) (Huang et al., 2009). The GO terms in biological processes and the KEGG pathways with *p* values < 0.005 and < 0.05, respectively, were defined as significantly enriched gene clusters. Scatter plots of the DEGs were visualized using R software (R Development Core Team) with a Bioconductor package, Reactome Pathway Analysis (Yu and He, 2015).

### STRING Analysis

Functional and physiological interactions of the iSN04-dependent DEGs were visualized using STRING version 11.0 (https://string-db.org/) (Szklarczyk et al., 2019).

### Basic Local Alignment Search Tool (BLAST) Search

Homologous sequences of iSN04 in the genomes of humans (taxid: 9605) and mice (taxid: 10088) were searched and scored using BLAST (https://blast.ncbi.nlm.nih.gov/Blast.cgi).

### Trivial Trajectory Parallelization of Multicanonical Molecular Dynamics (TTP-McMD)

Starting with the simulation of a single-chain iSN04 structure built from its DNA sequence by NAB in AmberTools (Macke and Case, 1998), enhanced ensemble method, TTP-McMD (Ikebe et al., 2011) was conducted, to sample the equilibrated conformations at 310 K. In the TTP-McMD, the energy range of the multicanonical ensemble covered a temperature range from 280 K to 380 K. Sixty trajectories were used and the production run was conducted for 40 ns in each trajectory (total 2.4 μs). Throughout the simulation, the force field of amber ff12SB (Maier et al., 2015) was used for iSN04, whereas the solvation effect was considered as a generalized-born model (Tsui and Case, 2001). The force field for the berberine molecule was constructed from the RESP charge assigned by the quantum mechanics result of the DFT method with B3-LYP/6-31G*, and the other parameters were taken from GAFF (Wang et al., 2004). In the initial structure of the iSN04-berberine system, a berberine molecule were put at a distance of 40 Å from iSN04. The conformation of the iSN04-berberine complex was calculated via TTP-McMD under the same conditions as the iSN04 simulation.

### Agarose Gel Electrophoresis

0.8 nmol PS-ODNs and 0.8 nmol berberine analogs were mixed in 16 μl Ham’s F10 medium (Supplementary Table S4). In the experiments shown in Supplementary Figure S5B, iSN04 and berberine were mixed in sterile water, or 4.5 mM of HCl, NaCl, MgCl_2_, KCl, CaCl_2_, or MnCl_2_ solution, or 0.45 mM of FeSO_4_, CuSO_4_, or ZnSO_4_ solution. The mixtures were placed at 4°C overnight, and then subjected to agarose gel electrophoresis using a TAE-buffered 3% agarose gel with 0.5 μg/ml ethidium bromide (EtBr). For colored images, the gels were illuminated by 302 nm ultraviolet (UV) using a UV Transilluminator (UVP, CA, USA) and the images were captured by a digital still camera without any filters. For monochromatic images, the gels were illuminated by 365-nm UV and the images were taken using ImageQuant LAS 500 with an emission bandpass filter of 560 nm (GE Healthcare).

### Protein Precipitation, SDSPAGE, and CBB Staining

Soluble whole-cell lysates of C2C12 and MC3T3-E1 cells were prepared using lysis buffer consisting of 0.1 M Tris-HCl (pH7.4), 75 mM NaCl, and 1% Triton X-100 (Nacalai) with protease inhibitor cocktail (1 mM 4-(2-aminoethyl)benzenesulfonyl fluoride hydrochloride, 0.8 μM aprotinin, 15 μM E-64, 20 μM leupeptin hemisulfate monohydrate, 50 μM bestatin, and 10 μM pepstatin A) (Nacalai). The biotin-conjugated PS-ODNs were immobilized on streptavidin-coated magnetic beads (Magnosphere MS300/Streptavidin; JSR Life Sciences, CA, USA) according to the manufacturer’s instruction. 100 μg of lysates and 0.6 mg of iSN14-beads were mixed in 1 ml lysis buffer with 1% NP-40 (Nacalai), and then gently rotated at 4°C overnight to eliminate the non-specific proteins absorbing onto ODNs or beads. After magnetic pull-down of iSN14-beads, the supernatants were admixed with iSN04-beads and rotated at 4°C overnight. The proteins precipitated by iSN04-beads were dissociated in lysis buffer with 1% NP-40, 10% glycerol, 2% sodium dodecyl sulfate (SDS) at 95°C for 5 min. The supernatants were subjected to SDS-PAGE using an 8% polyacrylamide gel. The gel was subjected to CBB staining using CBB Stain One Super (Nacalai) and scanned using ImageQuant LAS 500.

### Mass Spectrometry

The proteins within the CBB-stained gel were identified by mass spectrometry (MS Bioworks, MI, USA). In-gel digestion was performed using the ProGest robot (Digilab, MA, USA). The gels were washed with 25 mM ammonium bicarbonate followed by acetonitrile, reduced with 10 mM dithiothreitol at 60°C followed by alkylation with 50 mM iodoacetamide at room temperature, digested with trypsin (Promega) at 37°C for 4 h, and quenched with formic acid. Then the supernatant was subjected to analysis by nano LC-MS/MS with a Waters NanoAcquity HPLC system interfaced to a Thermo Fisher Q Exactive. Peptides were loaded on a trapping column and eluted over a 75-μm analytical column at 350 nl/min. Both columns were packed with Luna C18 resin (Phenomenex, CA, USA). The mass spectrometer was operated in data-dependent mode, with the Orbitrap operating at 70,000 FWHM and 17,500 FWHM for MS and MS/MS, respectively. The 15 most abundant ions were selected for MS/MS analysis. Data were searched using a local copy of Mascot with the following parameters: Enzyme, trypsin/P; Database, SwissProt Mouse; Fixed modification, carbamidomethyl; Variable modifications, oxidation, acetyl, pyro-Glu, deamidation; Mass values, monoisotopic; Peptide mass tolerance, 10 ppm; Fragment mass tolerance, 0.02 Da; Max missed cleavages, 2. Mascot DAT files were parsed into Scaffold (Proteome Software, OR, USA) for validation, filtering, and to create a non-redundant list per sample. Data were filtered using 1% protein and peptide FDR, which required at least two unique peptides per protein.

### Western Blotting

Soluble whole-cell lysates of the hMBs treated with 30 μM of iSN04 or AS1411 in DM for 48 h were prepared as described above. The lysates were denatured with 50 mM Tris-HCl, 10% glycerol, and 2% SDS at 95°C for 5 min. 10 μg of protein samples were subjected to SDS-PAGE on a 10% polyacrylamide gel followed by Western blotting using an iBlot 2 Dry Blotting System (Thermo Fisher Scientific). 1.0 μg/ml each of rabbit polyclonal anti-nucleolin antibody, mouse monoclonal anti-p53 antibody (PAb 240; Abcam), and mouse monoclonal anti-glyceraldehyde 3-phosphate dehydrogenase (GAPDH) antibody (5A12; Wako) were used as primary antibodies. 0.1 μg/ml each of horseradish peroxidase (HRP)-conjugated goat anti-rabbit and anti-mouse IgG antibodies (Jackson ImmunoResearch) were used as secondary antibodies, respectively. HRP activity was detected using ECL Prime reagents and ImageQuant LAS 500. The quantities of nucleolin and p53 proteins were normalized to that of GAPDH using ImageJ software.

### Statistical Analyses

Results are presented as the mean ± standard error. Statistical comparisons were performed using unpaired two-tailed Student’s *t* test, multiple comparison test with Dunnett’s test, Tukey-Kramer test, Scheffe’s *F* test, or Williams’ test where appropriate following one-way analysis of variance using R software. Statistical significance was set to *p* < 0.05.

## Results

### Identification of myoDNs

Fifty 18-base PS-ODNs (iSN01-iSN50) (Supplementary Table S1) derived from the LGG genome were subjected to a screening system to investigate the effects on myogenic differentiation of primary-cultured mMBs. 10 μM PS-ODNs were administered to the mMBs maintained in GM for 48 h. The mMBs were immunostained for MHC, a terminal differentiation marker of skeletal muscle (Supplementary Figure S1). The percentages of MHC^+^ cells were automatically quantified in a non-biased manner. As shown in Figure 1A, seven PS-ODNs (iSN01-iSN07) significantly increased the ratio of MHC^+^ cells, but other PS-ODNs did not alter the differentiation of mMBs. iSN01-iSN07 reproducibly induced myogenic differentiation of another independently isolated lot of mMBs (Supplementary Figure S2A), regardless of variation in the basal differentiation efficiency. In both screening results, iSN04’ exhibited the highest myogenetic activity (Figure 1A and Supplementary Figure S2A). These experiments were performed using iSN04’ (A<u>AG</u> TTA GGG TGA GGG TGA; not existing in LGG genome) instead of iSN04 (A<u>GA</u> TTA GGG TGA GGG TGA; existing in LGG genome). As the activities of iSN04’ and iSN04 were completely equal (Supplementary Figure S2C), iSN04 was utilized in the following experiments. iSN04 also promoted myogenic differentiation of the murine myoblast cell line C2C12 (Supplementary Figure S2D) and primary-cultured hMBs (Figure 1B). The ratio of MHC^+^ myocytes and fused myotubes was significantly increased by iSN04 in myoblasts in both mice and humans.

**Figure 1.**
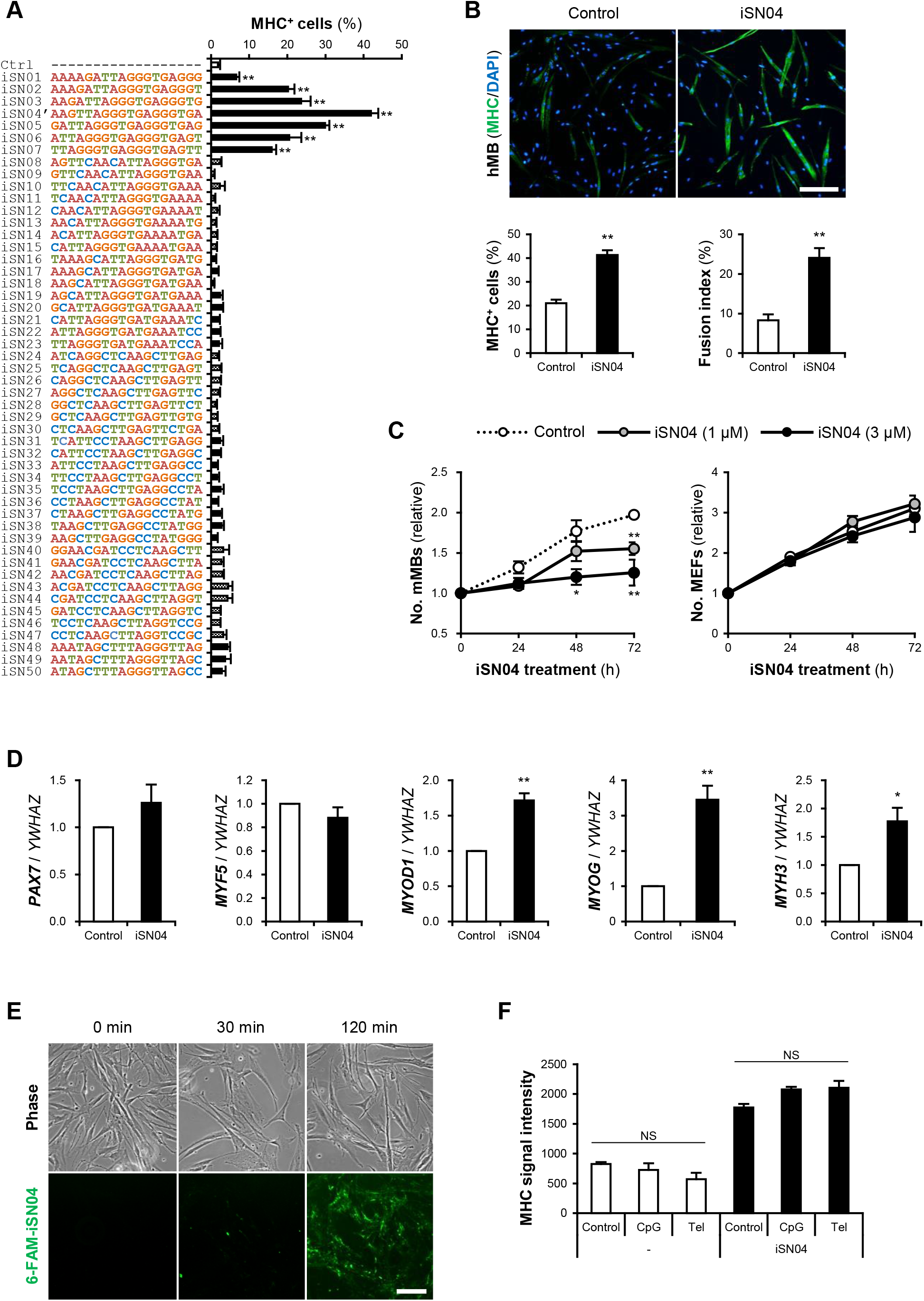
Identification of myoDNs. (**A**) Ratio of MHC^+^ cells within the screened mMBs treated with 10 μM PS-ODNs in GM for 48 h (screening system). ** *p* < 0.01 (Dunnett’s test). *n* = 3. (**B**) Representative immunofluorescent images of the hMBs treated with 10 μM iSN04 in DM for 48 h. Scale bar, 200 μm. Ratio of MHC^+^ cells and multinuclear myotubes were quantified. ** *p* < 0.01 (Student’s *t* test). *n* = 6. (**C**) Relative numbers of the mMBs and MEFs treated with 1 or 3 μM iSN04 in GM for each cell. Mean value of the control sample at 0 h was set to 1.0 for each experiment. * *p* < 0.05, ** *p* < 0.01 vs control at each time point (William’s test). *n* = 3. (**D**) qPCR results of myogenic gene expression in the hMBs treated with 30 μM iSN04 in DM for 24 h. Mean value of control hMBs was set to 1.0. * *p* < 0.05, ** *p* < 0.01 vs control (Student’s *t* test). *n* = 3. (**E**) Representative fluorescent images of the hMBs treated with 5 μg/ml 6-FAM-iSN04 in GM. Scale bar, 100 μm. (**F**) MHC signal intensities of the hMBs treated with 30 μM of iSN04, CpG-2006, or Tel-ODN in DM for 48 h (screening system). NS, no significant difference (Scheffe’s *F* test). *n* = 3.

In stem cells or their progenies, proliferation and differentiation are inverse processes, which negatively regulate each other (Ruijtenberg and van den Heuvel, 2016). The number of mMBs treated with iSN01-iSN07 was significantly lower than that of the control (Supplementary Figure S2B), indicating that iSN01-iSN07 inhibits myoblast proliferation. Continuous cell counting revealed that iSN04 suppressed the growth of mMBs in a dose-dependent manner; however, iSN04 did not alter the number of MEFs (Figure 1C). This demonstrates that the reduction in cell numbers in the iSN04-treated myoblasts was due to enhanced myogenic differentiation. qPCR revealed that iSN04 significantly upregulated the levels of myogenic transcription factors MyoD (*MYOD1*) and myogenin (*MYOG*), resulting in marked induction of embryonic MHC (*MYH3*) in hMBs (Figure 1D). In contrast, iSN04 did not alter the levels of undifferentiated myoblast markers, Pax7 (*PAX7*) and Myf5 (*MYF5*). These data show that iSN04 inherently promotes myoblast differentiation by activating the myogenic gene expression program.

We designated iSN01-iSN07 as “myoDNs”, denoting myogenetic ODNs. They are a novel type of ODNs that induce myoblast differentiation.

### myoDN Activity Is Independent of TLR Signaling

iSN01-iSN07 share a tandem repeat of a telomeric hexamer (TTAGGG TGAGGG) (Supplementary Figure S2E). A previous study has reported that a 24-base telomeric iODN (Tel-ODN) (Supplementary Table S1) suppresses human B-cell activation depending on TLR3, TLR7, and TLR9 (Sackesen et al., 2013). We have indicated that a 17-base telomeric iODN (iSG3; CCTCA TTAGGG TGAGGG) inhibits CpG-ODN (MsST)-induced interleukin (IL)-6 expression through intracellular incorporation in murine macrophages (Wang et al., 2015). Administration of 6-FAM-conjugated iSN04 to hMBs also showed that iSN04 was internalized into the cytoplasm within 2 h (Figure 1E). Contrarily, we have already confirmed that iSN01-iSN07 does not affect MsST-induced IL-6 expression in murine splenocytes (Nigar et al., 2017), suggesting that myoDNs are not iODNs. RT-PCR revealed that hMBs, mMBs, and C2C12 cells expressed all TLR genes except for *Tlr12* in C2C12 cells (Supplementary Figures S3A and S3B). To investigate the dependency of myoDN activity on TLR signaling, hMBs were treated with iSN04, Tel-ODN, or CpG-2006 (Supplementary Table S1). CpG-2006 is the TLR9 ligand initiating inflammatory responses in human lymphocytes and murine macrophages (Pohar et al., 2015). In the absence of iSN04, neither CpG-2006 nor Tel-ODN induced the differentiation of hMBs into MHC^+^ myocytes (Figure 1F). In the presence of iSN04, neither CpG-2006 nor Tel-ODN inhibited iSN04-induced myogenic differentiation. RNA-seq data (see next section) showed that transcription levels of the genes involved in the TLR signaling pathway were not altered in the iSN04-treated hMBs (Supplementary Figure S3C). These results demonstrate that iSN04 is not a TLR ligand, dissimilar to immunogenic CpG-ODNs and iODNs. It is assumed that myoDN activity inducing myogenic differentiation is independent of TLR signaling.

### Profile of iSN04-Dependent Gene Expression

We comprehensively surveyed the iSN04-dependent gene expression profile of hMBs. Total RNA of the hMBs treated with 30 μM iSN04 in DM for 24 h was subjected to RNA-seq (Supplementary Figure S4A). 51.3 million reads per sample were acquired, of which approximately 45.9 million reads (90.0%) were mapped to a human reference genome (Supplementary Table S3). In total, 60,448 transcripts were identified and their expression levels were calculated as FPKM. FPKM values of myogenic genes exhibited a pattern compatible with qPCR results; iSN04 significantly downregulated *MYF5* and upregulated *MYOD1* and *MYOG* (Supplementary Figure S4B). A total of 22,269 transcripts showed significant expression levels (FPKM > 0.1) in the control or iSN04 group. Of them, 899 transcripts were differentially expressed (> 1.5-fold) with the significance of FDR *p* < 0.05 (Supplementary Data). Of which 476 and 423 transcripts were upregulated and downregulated by iSN04, respectively. These DEGs depending on iSN04 were subjected to GO analysis. The 476 iSN04-upregulated DEGs significantly formed multiple gene clusters for muscle adaptation, contraction, and formation, which abundantly included sarcomeric components (myosin, actin, troponin, and their associated proteins) and transcription factors (myogenin, Hes1, Smad7, and Wnt10a) (Figure 2A). In contrast, the 423 iSN04-downregulated DEGs involved many clusters related to cell cycle and proliferation with higher significance (Figure 2B). These expression profiles of the iSN04-dependent DEGs corresponded well with the phenotype of iSN04-treated myoblasts, which showed promoted myogenic differentiation and arrested cell growth. STRING analysis visualized functional and physiological interactions of the DEGs or their products (Figure 2C). The tightly connected networks were detected in both DEG groups. Especially within the iSN04-downregulated group, 173 of the 423 DEGs (40.9%) were concentrated in the primary cluster, suggesting that iSN04 possibly suppresses at least one of the major nodes of the transcriptome at the early stage regulating myoblast fate. These data indicate that iSN04 globally modulates gene expression by orchestrating the myogenic program and cell cycle in myoblasts.

**Figure 2.**
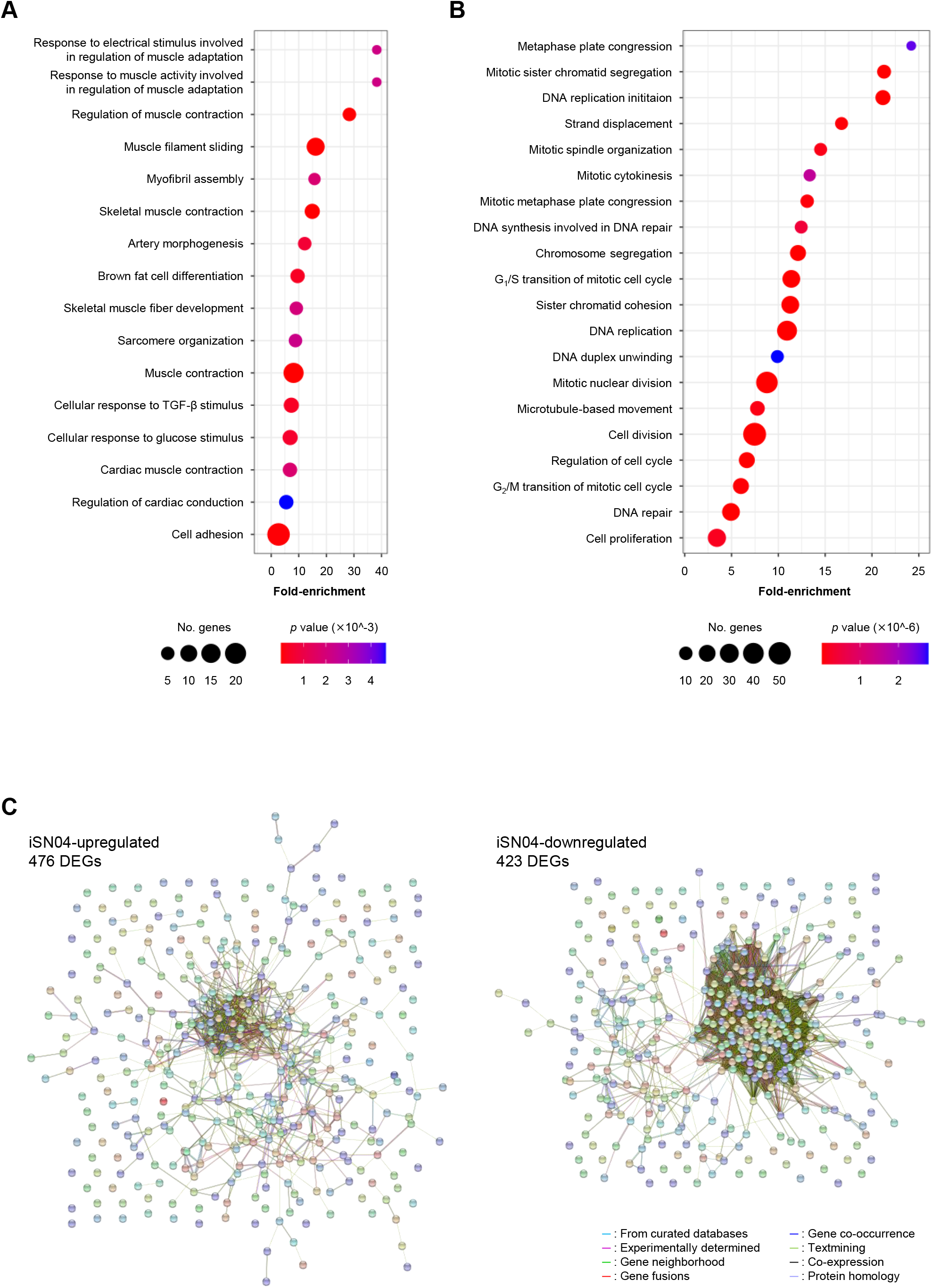
Profile of iSN04-dependent gene expression. (**A**) Scattered plot of the 476 iSN04-upregulated DEGs significantly (*p* < 5.0×10^-3^) enriched in GO terms. (**B**) Scattered plot of the 423 iSN04-downregulated DEGs significantly (*p* < 5.0×10^-6^) enriched in GO terms. (**C**) Functional and physiological networks within the 476 iSN04-upregulated DEGs (left panel) and the 423 iSN04-downregulated DEGs (right panel) visualized via STRING analysis.

### myoDN Activity Is Dependent on Its Structure

ODNs are classified into three categories according to their mechanism of action: antisense nucleotides, aptamers, and immunogenic ODNs as TLR ligands. myoDN activity was independent of TLR signals. Furthermore, immunogenic ODNs are often species-specific (Pohar et al., 2015), but iSN04 induces the differentiation of both murine and human myoblasts. To investigate the potential of myoDNs as antisense ODNs, the homologous sequences of iSN04 in human and murine genomes were surveyed using BLAST. The BLAST results displayed 59 loci in humans and 39 loci in mice that had iSN04-homologous sequences. However, there was no common gene or locus between humans and mice, denying that iSN04 serves as an antisense nucleotide. Intriguingly, the heat-denatured iSN02 lost the ability to induce myoblast differentiation (Supplementary Figure S2F), which strongly suggests that myoDN activity arises from its structure. Notably, iSN04 was resistant to thermal denaturation (Supplementary Figure S2G). The iSN04 conformation is considered to be relatively stable and can recover from denaturation in a short period. This might also be the reason why iSN04 presented the highest activity among the myoDNs.

The conformational properties of iSN04 under water conditions were computationally investigated using TTP-McMD (Ikebe et al., 2011). iSN04 at 310 K showed a compact globular structure (average radius: 0.96 nm), not a linear strand (Figure 3A). iSN04 displayed varied conformations, but their variations seemed to be limited within a certain range (Supplementary Movie). For fine conformation analysis, the contact probabilities between the residues of iSN04 were calculated. The ensemble-averaged contact probabilities at 310 K over all the simulated iSN04 structures were rendered as a contact map (Figure 3B). Three guanines at the 13-15th bases stacked upon each other, suggesting that this G_13-15_ stack is the stable center of the iSN04 structure. The impact of the G_13-15_ bases on iSN04 activity was examined using mutant iSN04. A series of deletions in the G_13-15_ bases gradually attenuated the myogenetic activity of iSN04. In particular, iSN04^Δ13^-^15^ completely lost its activity (Figure 3G), demonstrating that the G_13-15_ stack is indispensable for iSN04 activity.

**Figure 3.**
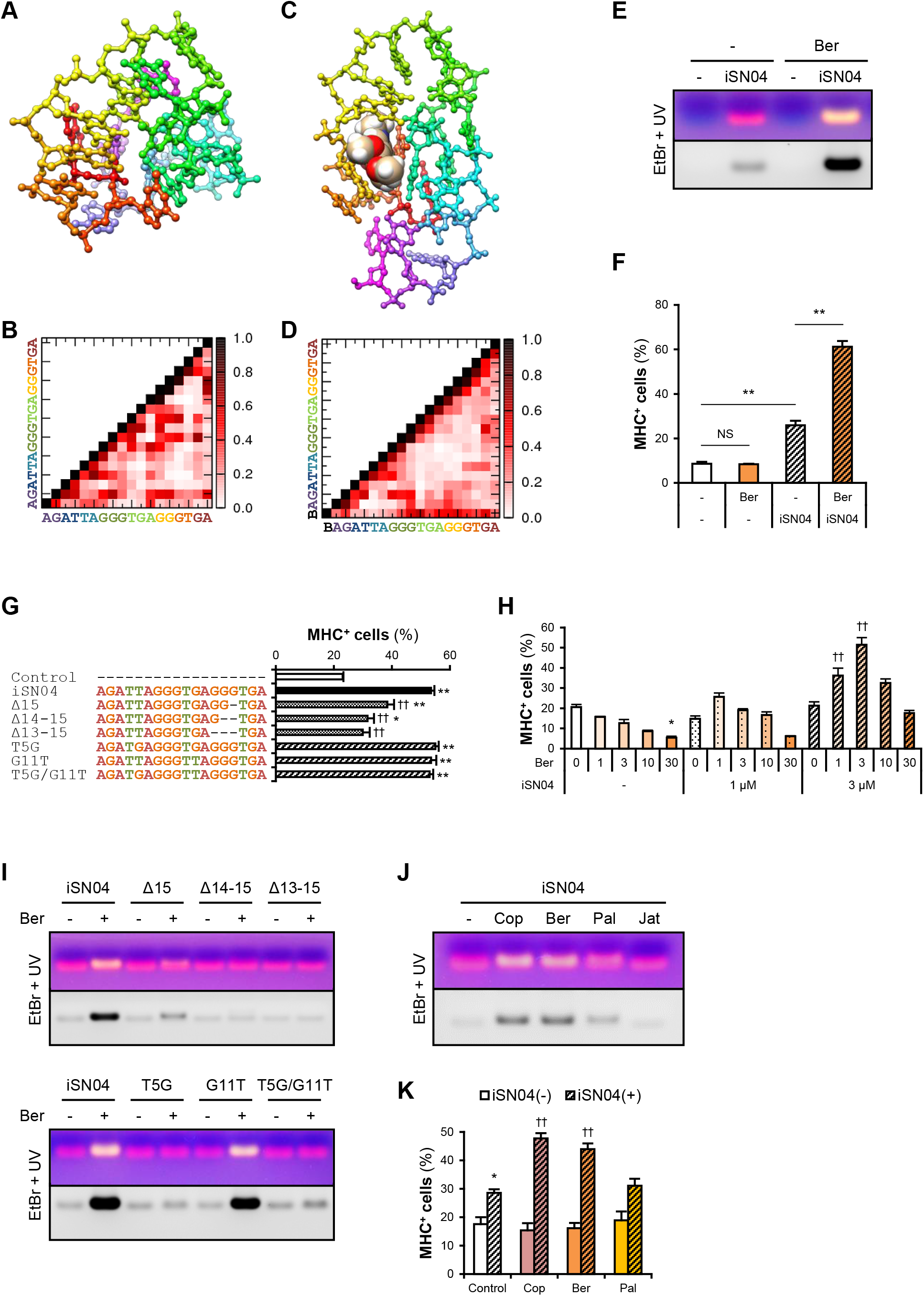
Berberine enhances iSN04 activity by causing a shift in molecular structure. (**A**) The conformation of iSN04 simulated via TTP-McMD. (**B**) Contact map of iSN04. The scale indicates contact probability. (**C**) The simulated conformation of iSN04-berberine complex. Berberine is shown as a sphere model. (**D**) Contact map of iSN04-berberine complex. “B” indicates berberine. (**E**) Representative images of agarose gel electrophoresis of iSN04 mixed with berberine (Ber) in F10 medium. The EtBr-stained and UV-irradiated gel was scanned without filters (upper image) and with a 560-nm filter (lower image). (**F**) Ratio of MHC^+^ cells within the mMBs treated with 10 μM iSN04 and 10 μM berberine in GM for 48 h (screening system). ** *p* < 0.01; NS, no significant difference (Scheffe’s *F* test). *n* = 3. (**G**) Ratio of MHC^+^ cells within the mMBs treated with 10 μM of mutant iSN04 in GM for 48 h (screening system). * *p* < 0.05, ** *p* < 0.01 vs control; ^‡‡^ *p* < 0.01 vs iSN04 (Scheffe’s *F* test). *n* = 3. (**H**) Ratio of MHC^+^ cells within the mMBs treated with 0, 1, or 3 μM iSN04 and 0, 1, 3, 10, or 30 μM of berberine in GM for 48 h (screening system). * *p* < 0.05 vs 0 μM-iSN04 + 0 μM-berberine; † *p* < 0.05, †† *p* < 0.01 vs 3 μM-iSN04 + 0 μM-berberine (Scheffe’s *F* test). *n* = 3. (**I**) Representative images of agarose gel electrophoresis of mutant iSN04 mixed with berberine in F10 medium. (**J**) Representative images of agarose gel electrophoresis of iSN04 mixed with coptisine (Cop), berberine, palmatine (Pal), or jatrorrhizine (Jat) in F10 medium. (**K**) Ratio of MHC^+^ cells within the mMBs treated with 10 μM iSN04 and 10 μM berberine analogs in GM for 48 h (screening system). * *p* < 0.05 vs control-iSN04(-); ^††^ *p* < 0.01 vs control-iSN04(+) (Tukey-Kramer test). *n* = 3.

### Berberine Enhances iSN04 Activity

We hypothesized that stabilization of the G_13-15_ stack would improve iSN04 activity. Telomeric DNA is known to form a highly ordered conformation. The G-quartet is a square aromatic surface constructed by the four guanines interacting with its neighbors via two hydrogen bonds. G-quartets stack upon each other to build the four-stranded G-quadruplex (Ou et al., 2008). Berberine, an isoquinoline alkaloid (Supplementary Figure S5A), interacts with the G-quartet and stabilizes the G-quadruplex structure derived from human telomeric DNA (Bazzicalupi et al., 2012). We tested whether iSN04 physically interacts with berberine. Briefly, iSN04 and berberine were mixed in F10 medium (Supplementary Table S4), subjected to electrophoresis, and stained with EtBr, which fluoresces red at 620 nm. The iSN04-berberine complex can be imaged with yellow fluorescence because berberine fluoresces green at 530 nm (Guo et al., 2015; Shinji et al., 2020). Indeed, yellow emissions of the iSN04-berberine complex were detected at a slightly higher molecular weight compared to iSN04 alone (Figure 3E). Not only iSN04 but also all other myoDNs interacted with berberine (Supplementary Figure S5C). G-quartets generally coordinate cations (Ou et al., 2008), and berberine binds to the telomeric DNA holding K^+^ (Bazzicalupi et al., 2012). We examined the requirement of cations for the iSN04-berberine complex using cationic solutions. Ca^2+^ was found to be necessary for the iSN04-berberine complex (Supplementary Figure S5B). Moreover, Mg^2+^ facilitated the interaction between iSN04 and berberine, but its effect was markedly weaker than that of Ca^2+^. These results suggest that berberine binds to the G-quartet-or G-quadruplex-like structure within iSN04, which is probably formed by the G_13-15_ stack.

Administration of iSN04 and berberine to mMBs proved that the activity of the iSN04-berberine complex was significantly higher than that of single iSN04 (Figure 3F). As berberine alone did not alter myoblast differentiation, it is possible that the improved activity of the iSN04-berberine complex is not a synergistic effect. Berberine is speculated to enhance the inherent activity of iSN04 by stabilizing or shifting the conformation. In some cases, one G-quartet binds to two berberine molecules (Bazzicalupi et al., 2012). To optimize the molar ratio of iSN04 to berberine, mMBs were treated with 0-3 μM iSN04 and 0-30 μM berberine. iSN04 exhibited the highest myogenetic activity when mMBs were co-treated with an equal molar of berberine (Figure 3H). Conformation of the iSN04-berberine complex at a molar ratio of 1:1 was simulated using TTP-McMD. Berberine interacted exactly with the G_13-15_ stack of iSN04 (Figure 3C). Deletions of the G_13-15_ bases of iSN04 experimentally demonstrated that berberine actually interacts with these guanines (Figure 3I, upper panel). The contact map of the iSN04-berberine complex showed that the G_7-9_ bases are stacked in addition to the G_13-15_ stack (Figure 3D). Berberine also contacted the G_9_ and consequently, it fits into the pocket assembled from the G_7-9_ and G_13-15_ stacks. iSN04 has two telomeric hexamers; T<u>T</u>AGGG and T<u>G</u>AGGG. We investigated the influence of the T_5_ and G_11_ of iSN04 on myogenetic activity and berberine binding. Both the T5G and G11T substitutions did not affect iSN04 activity (Figure 3G). However, the T5G substitution interfered with the formation of iSN04-berberine complex, and iSN04^G11T^ interacted with berberine as well as intact iSN04 (Figure 3I, lower panel). The contact maps indicated that the T_5_ relatively remains at a distance from other bases. On the contrary, guanines tend to interact with other bases. The extra guanine inserted by the T5G substitution might have perturbed the pocket required for iSN04 to bind to berberine.

We further examined the iSN04-enhancing abilities of three berberine analogs, coptisine, palmatine, and jatrorrhizine (Supplementary Figure S5A). Coptisine and berberine formed a complex with iSN04. Palmatine weakly interacted with iSN04, but jatrorrhizine did not interact at all (Figure 3J). Correspondingly, coptisine significantly improved the myogenetic activity of iSN04 to the same level as the iSN04-berberine complex (Figure 3K). These results illustrate that the 2,3-methylenedioxy ring of the berberine backbone is important for interacting with iSN04.

### iSN04 Targets Nucleolin and Increases p53 Protein

The structure-dependent myogenetic activity of iSN04 suggests the presence of iSN04-target proteins. We surveyed iSN04-binding proteins by precipitation assay. Biotin-conjugated iSN04 was immobilized on streptavidin-beads at the 5’ or 3’ end (iSN04-5’-Bio and iSN04-3’-Bio, respectively). Soluble whole-cell lysates were pre-pulled-down with iSN14-beads to eliminate the absorption of non-specific proteins onto ODNs or beads. After removing off-target proteins, the lysates were precipitated with iSN04-beads, followed to SDS-PAGE and CBB staining. Surprisingly, iSN04-binding protein was not detected in C2C12 cell lysates (data not shown). Next, the lysate of the murine osteoblast cell line MC3T3-E1 was prepared because iSN04 affected MC3T3-E1 differentiation (unpublished data). Both iSN04-5’-Bio and iSN04-3’-Bio precipitated the identical single protein with an expected size of 112 kDa, which was not precipitated by iSN14-beads or beads alone (Figure 4A). Mass spectrometry identified the iSN04-binding protein as nucleolin (Supplementary Tables S5 and S6). Although the molecular weight of nucleolin is 77 kDa, it is practically detected at 100-110 kDa because of the acidic amino acids in the N-terminal domain (Jia et al., 2017). Nucleolin is a multifunctional phosphoprotein located in the nucleolus, cytoplasm, and plasma membrane depending on the context of cellular processes such as gene expression, protein shuttling, cytokinesis, and apoptosis. Expression and subcellular localization of nucleolin are frequently abnormal in rapidly growing cells, typically cancers (Jia et al., 2017). A recent study reported that the amount and localization of nucleolin is involved in myogenic differentiation of C2C12 cells (Tang et al., 2017), but its precise mechanism is still unknown.

**Figure 4.**
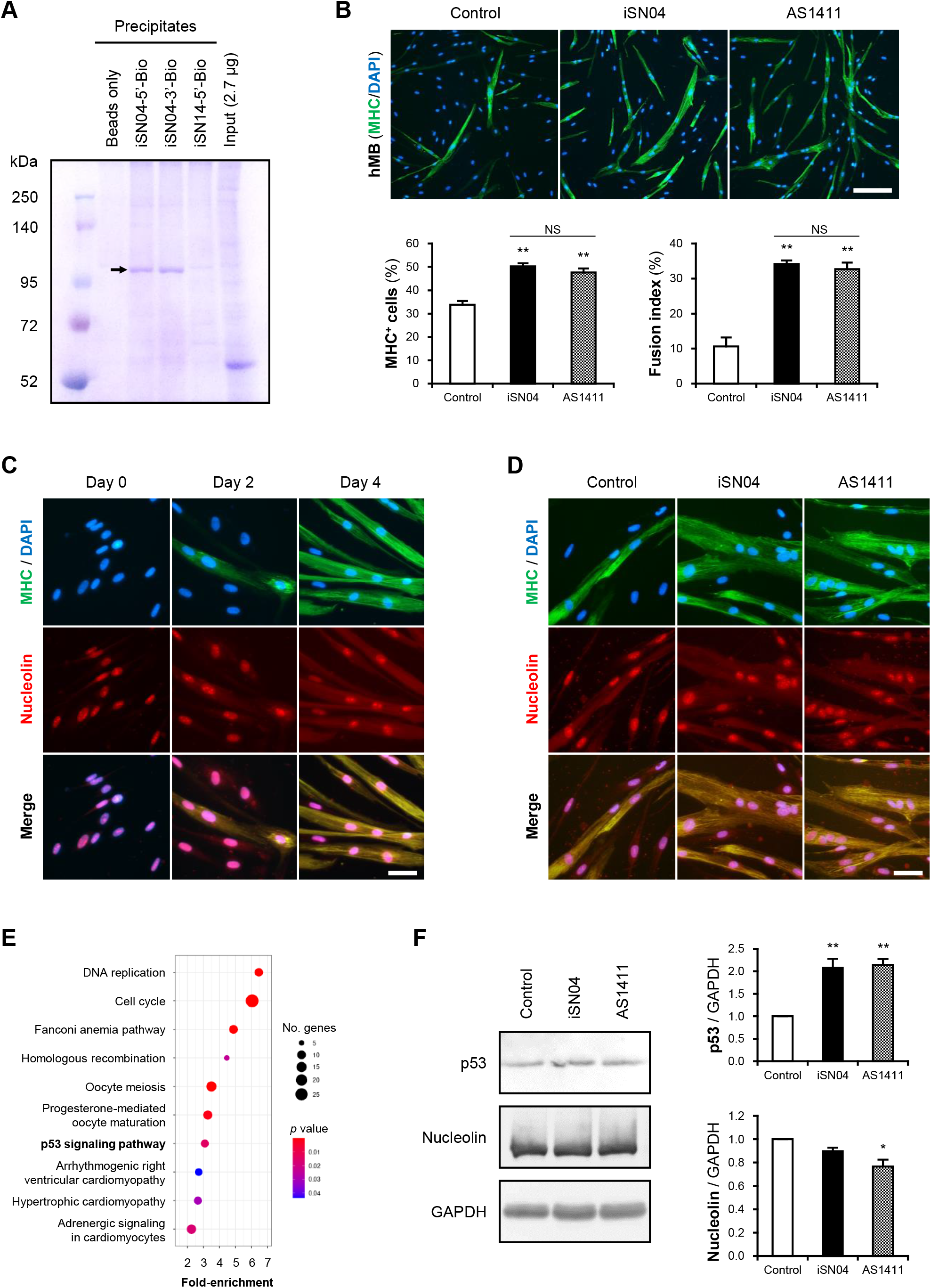
iSN04 targets nucleolin and improves p53 protein level. (**A**) Representative image of CBB-stained SDS-PAGE gel of the ODN-precipitated proteins. The arrow indicates the bands subjected to mass spectrometry. (**B**) Representative immunofluorescent images of the hMBs treated with 10 μM of iSN04 or AS1411 in DM for 48 h. Scale bar, 200 μm. Ratio of MHC^+^ cells and multinuclear myotubes were quantified. ** *p* < 0.01 vs control; NS, no significant difference (Student’s *t* test). *n* = 6. (**C**) Representative immunofluorescent images of the hMBs maintained in DM at day 0, 2, and 4. Scale bar, 50 μm. (**D**) Representative immunofluorescent images of the hMBs treated with 30 μM of iSN04 or AS1411 in DM for 48 h. Scale bar, 50 μm. (**E**) Scattered plot of the 899 iSN04-dependent DEGs significantly (FDR *p* < 0.05) enriched in KEGG pathways. (**F**) Representative images of Western blotting (10 μg protein/lane) of p53, nucleolin, and GAPDH in soluble whole cell lysates of the hMBs treated with 30 μM of iSN04 or AS1411 in DM for 48 h. Amounts of p53 and nucleolin were normalized using GAPDH. Mean value of control hMBs was set to 1.0. * *p* < 0.05, ** *p* < 0.01 vs control (Scheffe’s *F* test). *n* = 3.

A 26-base G-rich non-immunogenic ODN, AS1411 (Supplementary Table S1), is an established anti-nucleolin aptamer which has been utilized in several clinical trials on cancers (Bates et al., 2009; Yazdian-Robati et al., 2019). AS1411 promoted myogenic differentiation of hMBs to the same extent as iSN04 (Figure 4B), proving that iSN04 targets nucleolin in myoblasts. Immunostaining revealed that nucleolin initially localized in the nucleoli of hMBs and C2C12 cells in their undifferentiated states. Nucleolin was then translocated into the cytoplasm of MHC^+^ myotubes throughout myogenic differentiation (Figure 4C and Supplementary Figure S6A) as previously reported (Tang et al., 2017). The mRNA levels of nucleolin (*NCL*) in hMBs were increased during differentiation but did not chang after iSN04 treatment (Supplementary Figures S6C and S6D). In addition, nucleolin localization in hMBs or C2C12 cells was not shifted by iSN04 or AS1411 (Figure 4D and Supplementary Figure S6B). These results correspond to those of previous studies, which confirmed that AS1411 does not alter nucleolin localization in cancer cells (Litchfield et al., 2012; Reyes-Reyes et al., 2015; Ramos et al., 2020). Nucleolin has been reported to target the untranslated region (UTR) of p53 mRNA to interfere with its translation in tumor cells (Takagi et al., 2005; Chen et al., 2012). Nucleolin inhibition by AS1411 increases p53 protein levels and suppresses cell proliferation in glioma cells (Cheng et al., 2016). As p53 induces myogenic differentiation (Cerone et al., 2000; Porrello et al., 2000), antagonizing nucleolin in myoblasts by iSN04 or AS1411 was assumed to upregulate p53 protein and promote differentiation. RNA-seq data showed that the iSN04-dependent DEGs were enriched in the p53 signaling pathway (Figure 4E and Supplementary Figure S6E). Western blotting showed that both iSN04 and AS1411 increased p53 protein levels in hMBs (Figure 4F), even though iSN04 decreased the mRNA level of p53 (*TP53*) (Supplementary Figure S6F).

These data demonstrate that iSN04 antagonizes nucleolin to recover the suppressed p53 translation. The iSN04-increased p53 protein level activates downstream signal to arrest the cell cycle and induce myoblast differentiation. The results of this study present evidence that bacterial-derived ODNs can serve as aptamers to modulate cell fate.

## Discussion

To our knowledge, this is the first report of the ODNs promoting myogenic differentiation of skeletal muscle myoblasts. myoDNs are 18-base telomeric ODNs designed from the LGG genome sequence. Such bacterial ODNs serve as immunogenic ODNs recognized by TLRs and modulate the innate immune system (Krieg et al., 1995; Klinman, 2008). Among them, telomeric ODNs, also termed iODNs, are known to suppress inflammatory responses (Sackesen et al., 2013; Wang et al., 2015). In our previous study, myoDNs (iSN01-iSN07) were not iODNs (Nigar et al., 2017). iSN04, the myoDN presenting the highest activity, induced myoblast differentiation independent of TLR signaling. myoDNs are thus defined as a novel type of ODN that regulates cell fate through a unique mechanism. A previous study reported that CpG-2006 interferes with the osteoblastic differentiation of MSCs in a TLR9-independent manner, but its direct target is unknown (Norgaard et al., 2010). CpG-2006 was originally identified as a TLR9 ligand that activates immune responses (Hartmann et al., 2000; Bauer et al., 2001). The dual role of CpG-2006, in addition to myoDNs, implies that other bacterial ODNs might also exert non-immunological functions.

The present study revealed that the myogenetic activity of iSN04 arises from its conformation rather than its sequence. Molecular simulation and a series of mutant iSN04 demonstrated that the G_13-15_ stack within the second telomeric hexamer is essential for iSN04 activity. It is also indicated that berberine physically interacts with iSN04 via the G_13-15_ stack and enhances the myogenetic ability of iSN04, probably by stabilizing or optimizing the structure. This is the initial instance of functional improvement of ODNs by small molecules. Berberine is a safe isoquinoline alkaloid isolated from medicinal plants, that exhibits various bioactivities and has been utilized in clinical studies on diabetes, osteoporosis, and cancer (Imenshahidi and Hosseinzadeh, 2019). Berberine has also been studied as a ligand of the G-quadruplex, which is often formed in oncogenic promoter regions and regulates gene expression by recruiting transcriptional machinery (Siddiqui-Jain et al., 2002). Therefore, berberine derivatives that block G-quadruplexes can be potential anti-tumor drugs (Ou et al., 2008; Bazzicalupi et al., 2012). From these points of view, the modification of the structure and function of telomeric ODN by berberine provides a safe and reasonable technique for further research, development, and application of ODNs.

We identified nucleolin as a direct target of iSN04. The established anti-nucleolin aptamer, AS1411, also promoted myoblast differentiation, which proved that iSN04 antagonizes nucleolin. AS1411 has been reported to polymorphically fold into various conformations, including G-quadruplex structures (Dailey et al., 2010; Supplementary Figure S6G). Interestingly, iSN04 but not AS1411 interacted with berberine (Supplementary Figure S5D), and AS1411 but not iSN04 decreased nucleolin levels (Figure 4F), which suggests the presence of subtle structural and functional differences. Experimental determination of the iSN04 structure will provide valuable information on the similarity and dissimilarity between iSN04 and AS1411, which will be useful for building novel aptamers based on their conformations. Aptamers are usually developed via in vitro methodology using a specific target, systemic evolution of ligands by exponential enrichment (SELEX) (Wang et al., 2019). However, AS1411 is a non-SELEX aptamer that was screened as an inhibitor for cancer cell growth (Bates et al., 2009). Precipitation assay and mass spectrometry identified nucleolin as an AS1411 target (Girvan et al., 2006). Although AS1411 interacts with at least 15 proteins including nucleolin (Girvan et al., 2006), nucleolin has been a primary target of AS1411. Because aptamers usually recognize membrane proteins, and nucleolin is present on the plasma membrane of cancer cells (Bates et al., 2009; Jia et al., 2017; Yazdian-Robati et al., 2019).

Unlike many aptamers, iSN04 and AS1411 exert their effects inside myoblasts. iSN04 was spontaneously incorporated into myoblasts. Nucleolin initially localized in the nucleoli of growing myoblasts and diffused into the cytoplasm during myotube formation. However, iSN04 and nucleolin were not observed on the surface of myoblasts throughout differentiation. As discussed below, nuclear nucleolin serves as an mRNA-binding protein that regulates translation (Fahling et al., 2006). These findings indicate that iSN04 and AS1411 conceivably function in the nuclei of myoblasts. In general, single-strand ODNs are efficiently taken into the cytoplasm without carriers through gymnosis. Although its mechanism has not been completely understood, ODNs are considered to be incorporated by endocytosis, transported to the endosome, and are transferred to the cytoplasm through the endosomal membrane, probably due to their lower molecular weights and higher hydrophobicities compared to double-strand nucleotides. The released ODNs into the cytoplasm can accumulate in the nucleus by associating with chaperones or RNA-binding proteins (Juliano, 2018).

Nucleolin interferes with the translation of p53 mRNA by binding to its UTR (Takagi et al., 2005; Chen et al., 2012). Our study showed that antagonizing nucleolin by iSN04 or AS1411 increased p53 protein levels in myoblasts, as reported in AS1411-treated glioma cells (Cheng et al., 2016). The role of p53 in myoblasts has been intensively studied. An initial study found that the dominant-negative form of p53 inhibits the differentiation of C2C12 cells (Soddu et al., 1996). During myogenic differentiation, p53 cooperates with MyoD (Cerone et al., 2000) to activate transcription of retinoblastoma protein (Porrello et al., 2000), which serves as a cofactor of MyoD to arrest the cell cycle and facilitate muscle cell commitment (Gu et al., 1993; Novitch et al., 1996). A recent study revealed that p53 with MyoD coactivates the expression of the pro-apoptotic protein PUMA (Harford et al., 2017), which is required for the apoptosis associated with myoblast differentiation (Shaltouki et al., 2007; Harford et al., 2010). This accumulating evidence corroborates the findings that iSN04 upregulates p53 protein and induces myoblast differentiation.

Interestingly, iSN04 did not affect the growth of MEFs expressing nucleolin (Supplementary Figure S6H). In the precipitation assays, iSN04 pulled down nucleolin in the lysates of MC3T3-E1 cells but not of C2C12 cells, even though the amounts of nucleolin were nearly equal between the lysates (Supplementary Figure S6I). According to circumstances, nucleolin is post-translationally modified such as phosphorylation and glycosylation (Barel et al., 2001; Losfeld et al., 2009), and interacts with various partners including nucleotides and proteins (Jia et al., 2017). Probably due to that, AS1411 precipitates only certain forms of nucleolin (Teng et al., 2007; Bates et al., 2009). The amounts of the iSN04-binding form of nucleolin in C2C12 cells might be less than that in MC3T3-E1 cells and not enough to be detected via CBB staining. It is possible that the mode of existence of nucleolin differs among the cells, which affects its affinity to iSN04.

The precise role of nucleolin during myogenic differentiation is still not fully understood. A moderate decline in nucleolin protein by miR-34b has been reported to upregulate myogenic expression (Tang et al., 2017). This study showed that nucleolin levels decreased through differentiation of C2C12 cells; however, our results using primary-cultured hMBs showed increased nucleolin expression upon differentiation. As nucleolin is potently induced in actively proliferating cells like tumors (Jia et al., 2017), nucleolin levels might be high in the immortalized C2C12 cell line. Therefore, nucleolin function in myoblasts needs to be further investigated using primary-cultured cells or in vivo models. In both hMBs and C2C12 cells, nucleolin initially localized in the nucleoli and then diffused into the cytoplasm through differentiation. An analogous shift of nucleolin localization has been observed during adipogenic differentiation of 3T3-L1 pre-adipocytes (Wang et al., 2015). The biological activities of nucleolin can vary depending on its subcellular distribution. Numerous studies have revealed that nucleolar nucleolin regulates RNA metabolism, nucleoplasmic nucleolin modulates gene expression, cytoplasmic nucleolin serves as a shuttle protein, and cell surface nucleolin is involved in various signaling pathways (Jia et al., 2017). Elucidating the relationship between nucleolin localization and differentiation of precursor cells would be important to understand the fine-tuned mechanism of myoDNs.

To establish myoDNs as potential drug seeds for muscle diseases, their pharmacological actions need to be investigated in vivo. Intramuscular injection (i.m.) of antisense nucleotides have been clinically applied to treat Duchenne muscular dystrophy (Quemener et al., 2020). However, i.m. is painful and there is a risk of sterile abscess formation, therefore it is not suitable for prevention or treatment of the long-termed muscle atrophy associating with aging or chronic diseases. We have previously developed ODN nanocapsules (ODNcaps) as an oral delivery system of ODNs. Oral administration of the capsuled iODN to atopic model mice successfully suppressed immune responses in dermatitis (Wang et al., 2015). ODNcaps would be also useful to deliver myoDNs to atrophic muscle tissue. Further studies using adequate animal model and drug delivery system are required for clinical application of myoDNs in future.

## Conclusion

This study presents that bacterial genome-derived myoDNs promote myogenic differentiation by targeting nucleolin. The myoDN activities can be enhanced by conformational changes via binding to berberine. myoDNs are expected to be novel and unique drug candidates for muscle diseases, including atrophy, in which myoblasts are functionally deteriorated.

## Data Availability Statement

FASTQ raw read data of RNA-seq were deposited in the DDBJ Sequence Read Archive (DRA; Research Organization of Information and Systems, National Institute of Genetics, Mishima, Japan) with the accession number: DRA008498.

## Ethics Statement

All experimental procedures were conducted in accordance with the Regulations for Animal Experimentation of Shinshu University, and the animal experimentation protocol was approved by the Committee for Animal Experiments of Shinshu University.

## Author Contributions

TT designed the study; TT and KU wrote the manuscript; SS, YN, SN, and TT performed the experiments and data analyses; KU performed molecular simulation and proposed iSN04-berberine interaction; TS designed and provided the ODNs.

## Conflict of Interst

Shinshu University has been assigned the invention of myoDNs by TT, KU, and TS, and Japan Patent Application 2018-568609 has been filed on February 15, 2018.

## Acknowledgments

C57BL/6J mice and C2C12 cells were kindly provided by Dr. Sachi Tanaka and Dr. Shinichi Yonekura, of Shinshu University. The preprint has been posted on bioRxiv (doi: 10.1101/2020.10.07.330472).

