## Supplementary Information for "Identification of the myogenetic oligodeoxynucleotides (myoDNs) that promote differentiation of skeletal muscle myoblasts by targeting nucleolin" (from another version of this preprint: version 1, 2020-10-07)

**Table S1.** Sequences of PS-ODNs.

| Name | Sequence (5'-3') | Reference |
|---|---|---|
| iSN01 | AAAAGATTAGGGTGAGGG | Nigar et al., 2017 |
| iSN02 | AAAGATTAGGGTGAGGGT | Nigar et al., 2017 |
| iSN03 | AAGATTAGGGTGAGGGTG | Nigar et al., 2017 |
| iSN04 | AGATTAGGGTGAGGGTGA | Nigar et al., 2017 |
| iSN04' | AAGTTAGGGTGAGGGTGA | This study |
| iSN05 | GATTAGGGTGAGGGTGAG | Nigar et al., 2017 |
| iSN06 | ATTAGGGTGAGGGTGAGT | Nigar et al., 2017 |
| iSN07 | TTAGGGTGAGGGTGAGTT | Nigar et al., 2017 |
| iSN08 | AGTTCAACATTAGGGTGA | Nigar et al., 2017 |
| iSN09 | GTTCAACATTAGGGTGAA | Nigar et al., 2017 |
| iSN10 | TTCAACATTAGGGTGAAA | Nigar et al., 2017 |
| iSN11 | TCAACATTAGGGTGAAAA | Nigar et al., 2017 |
| iSN12 | CAACATTAGGGTGAAAAT | Nigar et al., 2017 |
| iSN13 | AACATTAGGGTGAAAATG | Nigar et al., 2017 |
| iSN14 | ACATTAGGGTGAAAATGA | Nigar et al., 2017 |
| iSN15 | CATTAGGGTGAAAATGAA | Nigar et al., 2017 |
| iSN16 | TAAAGCATTAGGGTGATG | Nigar et al., 2017 |
| iSN17 | AAAGCATTAGGGTGATGA | Nigar et al., 2017 |
| iSN18 | AAGCATTAGGGTGATGAA | Nigar et al., 2017 |
| iSN19 | AGCATTAGGGTGATGAAA | Nigar et al., 2017 |
| iSN20 | GCATTAGGGTGATGAAAT | Nigar et al., 2017 |
| iSN21 | CATTAGGGTGATGAAATC | Nigar et al., 2017 |
| iSN22 | ATTAGGGTGATGAAATCC | Nigar et al., 2017 |
| iSN23 | TTAGGGTGATGAAATCCA | Nigar et al., 2017 |
| iSN24 | ATCAGGCTCAAGCTTGAG | Nigar et al., 2017 |
| iSN25 | TCAGGCTCAAGCTTGAGT | Nigar et al., 2017 |
| iSN26 | CAGGCTCAAGCTTGAGTT | Nigar et al., 2017 |
| iSN27 | AGGCTCAAGCTTGAGTTC | Nigar et al., 2017 |
| iSN28 | GGCTCAAGCTTGAGTTCT | Nigar et al., 2017 |
| iSN29 | GCTCAAGCTTGAGTTGTG | Nigar et al., 2017 |
| iSN30 | CTCAAGCTTGAGTTCTGA | Nigar et al., 2017 |
| iSN31 | TCATTCCTAAGCTTGAGG | Nigar et al., 2017 |
| iSN32 | CATTCCTAAGCTTGAGGC | Nigar et al., 2017 |

| | | |
|---|---|---|
| iSN33 | ATTCCTAAGCTTGAGGCC | Nigar et al., 2017 |
| iSN34 | TTCCTAAGCTTGAGGCCT | Nigar et al., 2017 |
| iSN35 | TCCTAAGCTTGAGGCCTA | Nigar et al., 2017 |
| iSN36 | CCTAAGCTTGAGGCCTAT | Nigar et al., 2017 |
| iSN37 | CTAAGCTTGAGGCCTATG | Nigar et al., 2017 |
| iSN38 | TAAGCTTGAGGCCTATGG | Nigar et al., 2017 |
| iSN39 | AAGCTTGAGGCCTATGGG | Nigar et al., 2017 |
| iSN40 | GGAACGATCCTCAAGCTT | Nigar et al., 2017 |
| iSN41 | GAACGATCCTCAAGCTTA | Nigar et al., 2017 |
| iSN42 | AACGATCCTCAAGCTTAG | Nigar et al., 2017 |
| iSN43 | ACGATCCTCAAGCTTAGG | Nigar et al., 2017 |
| iSN44 | CGATCCTCAAGCTTAGGT | Nigar et al., 2017 |
| iSN45 | GATCCTCAAGCTTAGGTC | Nigar et al., 2017 |
| iSN46 | TCCTCAAGCTTAGGTCCG | Nigar et al., 2017 |
| iSN47 | CCTCAAGCTTAGGTCCGC | Nigar et al., 2017 |
| iSN48 | AAATAGCTTTAGGGTTAG | Nigar et al., 2017 |
| iSN49 | AATAGCTTTAGGGTTAGC | Nigar et al., 2017 |
| iSN50 | ATAGCTTTAGGGTTAGCC | Nigar et al., 2017 |
| iSN04$^{\Delta 15}$ | AGATTAGGGTGAGGTGA | This study |
| iSN04$^{\Delta 14\text{-}15}$ | AGATTAGGGTGAGTGA | This study |
| iSN04$^{\Delta 13\text{-}15}$ | AGATTAGGGTGATGA | This study |
| iSN04$^{T5G}$ | AGATGAGGGTGAGGGTGA | This study |
| iSN04$^{G11T}$ | AGATTAGGGTTAGGGTGA | This study |
| iSN04$^{T5G/G11T}$ | AGATGAGGGTTAGGGTGA | This study |
| CpG-2006 | TCGTCGTTTTGTCGTTTTGTCGTT | Pohar et al., 2015 |
| Tel-ODN | TTAGGGTTAGGGTTAGGGTTAGGG | Sackesen et al., 2013 |
| AS1411* | GGTGGTGGTGGTTGTGGTGGTGGTGG | Girvan et al., 2006 |

* AS1411 has a phosphodiester backbone. Other ODNs were PS-ODNs.

**Table S2.** Primer sequences for qPCR.

| Gene | Sequence (5'-3') | Size (bp) | Reference |
|---|---|---|---|
| *MYF5* | AGAACTACTATAGCCTGCCGG<br>ATCTGTGGCATATACATTTGATACATCA | 164 | Zibat et al., 2010 |
| *MYH3* | GGACAGGAAGAATGTGCTGAGATT<br>GCCTCTTGTAGGACTTGACTTTCAC | 74 | Norman et al., 2009 |
| *MYOD1* | TGCTCCGACGGCATGATGGAC<br>TCGACACCGCCGCACTCT | 127 | Sharma et al., 2013 |
| *MYOG* | AACCCAGGGGATCATCTGCTCAC<br>GTTGGGCATGGTTTCATCTGGGAAG | 129 | Sharma et al., 2013 |
| *NCL* | ATTGGTAGCAACTCCTGGTAAG<br>CACTGTCATCATCCTCCTCTTC | 110 | Cai et al., 2018 |
| *PAX7* | GACCCCTGCCTAACCACATC<br>GTCTCCTGGTAGCGGCAAAG | 133 | Sakai et al., 2017 |
| *TLR1* | CTATACACCAAGTTGTCAGC<br>GTCTCCAACTCAGTAAGGTG | 220 | Yao et al., 2015 |
| *TLR2* | GGCCAGCAAATTACCTGTGTG<br>AGGCGGACATCCTGAACCT | 67 | Hayashi et al., 2003 |
| *TLR3* | TCCCAAGCCTTCAACGACTG<br>TGGTGAAGGAGAGCTATCCACA | 68 | Hayashi et al., 2003 |
| *TLR4* | CTGCAATGGATCAAGGACCA<br>TTATCTGAAGGTGTTGCACATTCC | 74 | Hayashi et al., 2003 |
| *TLR5* | TCGAGCCCCTACAAGGGAA<br>CACTGAGACTCTGCTATACAAGCTA | 74 | Hayashi et al., 2003 |
| *TLR6* | CTATTGTTAAAAGCTTCCATTTTGT<br>ACCTGAAGCTCAGCGATGTAGTTC | 187 | Hayashi et al., 2003 |
| *TLR7* | TTACCTGGATGGAAACCAGCTAC<br>TCAAGGCTGAGAAGCTGTAAGCTA | 72 | Hayashi et al., 2003 |
| *TLR8* | GAGAGCCGAGACAAAAACGTTC<br>TGTCGATGATGGCCAATCC | 73 | Hayashi et al., 2003 |
| *TLR9* | TGGTGTTGAAGGACAGTTCTCTC<br>CACTCGGAGGTTTCCCAGC | 71 | Hayashi et al., 2003 |
| *TLR10* | GAAAGGTTCCCGCAGACTTG<br>TGGAGTTGAAAAAGGAGGTTATAG | 73 | Hayashi et al., 2003 |

| | | | |
|---|---|---|---|
| *YWHAZ* | CAAGCATACCAAGAAGCATTTGA<br>GGGCCAGACCCAGTCTGA | 76 | Nihashi et al., 2019 |
| *Tlr1* | TCTCTGAAGGCTTTGTCGATACA<br>GACAGAGCCTGTAAGCATATTCG | 212 | Caramalho et al., 2003 |
| *Tlr2* | TCTAAAGTCGATCCGCGACAT<br>TACCCAGCTCGCTCACTACGT | 344 | Caramalho et al., 2003 |
| *Tlr3* | TTGTCTTCTGCACGAACCTG<br>CGCAACGCAAGGATTTTATT | 205 | Caramalho et al., 2003 |
| *Tlr4* | CAAGAACATAGATCTGAGCTTCAACCC<br>GCTGTCCAATAGGGAAGCTTTCTAGAG | 278 | Caramalho et al., 2003 |
| *Tlr5* | ACTGAATTCCTTAAGCGACGTA<br>AGAAGATAAAGCCGTGCGAAA | 401 | Caramalho et al., 2003 |
| *Tlr6* | AACAGGATACGGAGCCTTGA<br>CCAGGAAAGTCAGCTTCGTC | 199 | Caramalho et al., 2003 |
| *Tlr7* | TTCCGATACGATGAATATGCACG<br>TGAGTTTGTCCAGAAGCCGTAAT | 412 | Caramalho et al., 2003 |
| *Tlr8* | GGCACAACTCCCTTGTGATT<br>CATTTGGGTGCTGTTGTTTG | 195 | Caramalho et al., 2003 |
| *Tlr9* | TGCAATTGGCTGTTCCTGAA<br>GGTGGTGGATACGGTTGGAG | 100 | Saluja et al., 2012 |
| *Tlr11* | CCAGGACTGCACCTTTTGG<br>GTGACACTGGTTGTACGCAAT | 185 | This study |
| *Tlr12* | AGAGCTGGCTGGTATGTTCC<br>GTGTTCTTGTCAGGTCCAGAATC | 161 | This study |
| *Tlr13* | GGAGCGCCTTGATCTAACTAACA<br>TCAGGTGGGTCAGAGAAACCA | 80 | This study |

**Table S3.** Numbers of the reads obtained by RNA-seq.

| Sample | Raw reads | Cleaned reads | Mapped reads | % mapped |
|---|---|---|---|---|
| Control-A | 59,865,454 | 58,169,640 | 53,874,761 | 92.3% |
| Control-B | 51,357,760 | 50,323,502 | 46,717,465 | 92.8% |
| Control-C | 46,217,046 | 42,965,166 | 39,340,050 | 91.6% |
| iSN04-A | 47,556,514 | 46,737,876 | 43,434,950 | 92.9% |
| iSN04-B | 45,959,342 | 45,008,402 | 41,670,855 | 92.6% |
| iSN04-C | 56,638,202 | 54,726,942 | 50,462,497 | 92.2% |

**Table S4.** Ionic components of Ham's F10 medium.

| Components | Concentration |
|---|---|
| $CaCl_2$ | 0.3 mM |
| $CuSO_4$ | $1\times10^{-5}$ mM |
| $FeSO_4$ | $1\times10^{-3}$ mM |
| $MgSO_4$ | 0.6 mM |
| KCl | 3.8 mM |
| $KH_2PO_4$ | 0.6 mM |
| $NaHCO_3$ | 14.3 mM |
| NaCl | 127.6 mM |
| $Na_2HPO_4$ | 1.1 mM |
| $ZnSO_4$ | $1\times10^{-4}$ mM |

**Table S5.** Summary of mass spectrometry.

| | **iSN04-5'-bio** | **iSN04-3'-bio** |
|---|---:|---:|
| No. of proteins identified | 14 | 74 |
| No. of unique peptides | 98 | 521 |
| No. of spectra matching | 116 | 715 |

**Table S6.** Results of mass spectrometry for iSN04-binding protein.

| Protein | Accession No. | Molecular weight | iSN04-5’-bio | iSN04-3’-bio |
|---|---|---|---|---|
| Nucleolin | P09405, NUCL_MOUSE | 77 kDa | 42 | 132 |
| Heat shock protein HSP 90-beta | P11499, HS90B_MOUSE | 83 kDa | 4 | 41 |
| DNA topoisomerase 1 | Q04750, TOP1_MOUSE | 91 kDa | 10 | 37 |
| Heat shock protein HSP 90-alpha | P07901, HS90A_MOUSE | 85 kDa | 0 | 36 |
| Eukaryotic translation initiation factor 4 gamma 2 | Q62448, IF4G2_MOUSE | 102 kDa | 0 | 20 |
| Elongation factor 2 | P58252, EF2_MOUSE | 95 kDa | 0 | 18 |
| Junctophilin-2 | Q9ET78, JPH2_MOUSE | 75 kDa | 0 | 18 |
| Nucleolar transcription factor 1 | P25976, UBF1_MOUSE | 90 kDa | 0 | 13 |
| 40S ribosomal protein S3 | P62908, RS3_MOUSE | 27 kDa | 0 | 10 |
| Transitional endoplasmic reticulum ATPase | Q01853, TERA_MOUSE | 89 kDa | 0 | 10 |
| Cyclin-dependent kinase 11B | P24788, CD11B_MOUSE | 92 kDa | 0 | 8 |
| Aspartyl/asparaginyl beta-hydroxylase | Q8BSY0, ASPH_MOUSE | 83 kDa | 0 | 7 |
| Splicing factor, proline- and glutamine-rich | Q8VIJ6, SFPQ_MOUSE | 75 kDa | 2 | 7 |
| Plasminogen activator inhibitor 1 RNA-binding protein | Q9CY58, PAIRB_MOUSE | 45 kDa | 2 | 7 |
| AP-2 complex subunit beta | Q9DBG3, AP2B1_MOUSE | 105 kDa | 0 | 7 |
| AP-2 complex subunit alpha-2 | P17427, AP2A2_MOUSE | 104 kDa | 0 | 6 |
| 40S ribosomal protein S2 | P25444, RS2_MOUSE | 31 kDa | 0 | 6 |
| 60S ribosomal protein L8 | P62918, RL8_MOUSE | 28 kDa | 0 | 6 |
| Epidermal growth factor receptor kinase substrate 8 | Q08509, EPS8_MOUSE | 92 kDa | 0 | 6 |

| | | | | |
|---|---|---|---|---|
| tRNA (cytosine(34)-C(5))-methyltransferase | Q1HFZ0, NSUN2_MOUSE | 85 kDa | 0 | 6 |
| Eukaryotic translation initiation factor 3 subunit B | Q8JZQ9, EIF3B_MOUSE | 91 kDa | 0 | 6 |
| 40S ribosomal protein S9 | Q6ZWN5, RS9_MOUSE | 23 kDa | 0 | 5 |
| Mannosyl-oligosaccharide glucosidase | Q80UM7, MOGS_MOUSE | 92 kDa | 0 | 5 |
| Desmoplakin | E9Q557, DESP_MOUSE | 333 kDa | 0 | 4 |
| Elongation factor 1-alpha 1 | P10126, EF1A1_MOUSE | 50 kDa | 0 | 4 |
| Putative ATP-dependent RNA helicase Pl10 | P16381, DDX3L_MOUSE | 73 kDa | 0 | 4 |
| ADP/ATP translocase 1 | P48962, ADT1_MOUSE | 33 kDa | 0 | 4 |
| DNA replication licensing factor MCM5 | P49718, MCM5_MOUSE | 82 kDa | 0 | 4 |
| Actin, cytoplasmic 1 | P60710, ACTB_MOUSE | 42 kDa | 0 | 4 |
| 40S ribosomal protein S6 | P62754, RS6_MOUSE | 29 kDa | 0 | 4 |
| Nuclear receptor subfamily 4 group A member 2 | Q06219, NR4A2_MOUSE | 67 kDa | 5 | 4 |
| Probable ATP-dependent RNA helicase DDX17 | Q501J6, DDX17_MOUSE | 72 kDa | 0 | 4 |
| Eukaryotic translation initiation factor 4B | Q8BGD9, IF4B_MOUSE | 69 kDa | 0 | 4 |
| 26S proteasome non-ATPase regulatory subunit 2 | Q8VDM4, PSMD2_MOUSE | 100 kDa | 0 | 4 |
| ATP-dependent RNA helicase DDX1 | Q91VR5, DDX1_MOUSE | 83 kDa | 0 | 4 |
| Polypyrimidine tract-binding protein 1 | P17225, PTBP1_MOUSE | 56 kDa | 0 | 3 |
| 60S ribosomal protein L3 | P27659, RL3_MOUSE | 46 kDa | 0 | 3 |
| 60S ribosomal protein L13 | P47963, RL13_MOUSE | 24 kDa | 0 | 3 |
| 60S ribosomal protein L9 | P51410, RL9_MOUSE | 22 kDa | 0 | 3 |
| 40S ribosomal protein S25 | P62852, RS25_MOUSE | 14 kDa | 0 | 3 |
| 60S ribosomal protein L10-like | P86048, RL10L_MOUSE | 25 kDa | 0 | 3 |

| | | | | |
|---|---|---|---|---|
| 40S ribosomal protein S3a | P97351, RS3A_MOUSE | 30 kDa | 0 | 3 |
| Lymphocyte-specific helicase | Q60848, HELLS_MOUSE | 95 kDa | 0 | 3 |
| Cytoskeleton-associated protein 4 | Q8BMK4, CKAP4_MOUSE | 64 kDa | 0 | 3 |
| Elongin-A | Q8CB77, ELOA1_MOUSE | 87 kDa | 0 | 3 |
| Ras and Rab interactor 1 | Q921Q7, RIN1_MOUSE | 83 kDa | 0 | 3 |
| Inversin | O89019, INVS_MOUSE | 117 kDa | 0 | 2 |
| Vitronectin | P29788, VTNC_MOUSE | 55 kDa | 0 | 2 |
| 40S ribosomal protein S20 | P60867, RS20_MOUSE | 13 kDa | 0 | 2 |
| Heterogeneous nuclear ribonucleoprotein K | P61979, HNRPK_MOUSE | 51 kDa | 0 | 2 |
| 40S ribosomal protein S14 | P62264, RS14_MOUSE | 16 kDa | 0 | 2 |
| 40S ribosomal protein S11 | P62281, RS11_MOUSE | 18 kDa | 0 | 2 |
| 40S ribosomal protein S4, X isoform | P62702, RS4X_MOUSE | 30 kDa | 0 | 2 |
| 60S ribosomal protein L22 | P67984, RL22_MOUSE | 15 kDa | 0 | 2 |
| 40S ribosomal protein S5 | P97461, RS5_MOUSE | 23 kDa | 0 | 2 |
| Mitogen-activated protein kinase kinase kinase kinase 5 | Q8BPM2, M4K5_MOUSE | 95 kDa | 0 | 2 |
| La-related protein 4 | Q8BWW4, LARP4_MOUSE | 80 kDa | 0 | 2 |
| NFX1-type zinc finger-containing protein 1 | Q8R151, ZNFX1_MOUSE | 219 kDa | 0 | 2 |
| Activating signal cointegrator 1 complex subunit 2 | Q91WR3, ASCC2_MOUSE | 86 kDa | 0 | 2 |
| Probable ATP-dependent RNA helicase DDX27 | Q921N6, DDX27_MOUSE | 86 kDa | 0 | 2 |
| Mitogen-activated protein kinase kinase kinase kinase 3 | Q99JP0, M4K3_MOUSE | 101 kDa | 0 | 2 |
| 60S ribosomal protein L15 | Q9CZM2, RL15_MOUSE | 24 kDa | 0 | 2 |
| Elongation factor 1-gamma | Q9D8N0, EF1G_MOUSE | 50 kDa | 0 | 2 |

| | | | | |
|---|---|---|---|---|
| N-terminal kinase-like protein | Q9EQC5, SCYL1_MOUSE | 89 kDa | 0 | 2 |
| Ribosomal protein S6 kinase alpha-4 | Q9Z2B9, KS6A4_MOUSE | 86 kDa | 0 | 2 |
| Stress-70 protein, mitochondrial | P38647, GRP75_MOUSE | 73 kDa | 2 | 0 |
| Adenosylhomocysteinase | P50247, SAHH_MOUSE | 48 kDa | 3 | 0 |
| ATP synthase subunit alpha, mitochondrial | Q03265, ATPA_MOUSE | 60 kDa | 2 | 0 |

Numerals are the number of the identified proteins in the samples.

### Figure Legends

**Figure S1.** Screening of the LGG-ODNs promoting myoblast differentiation. Representative immunofluorescent images of the mMBs treated with 10 µM PS-ODNs in GM for 48 h (screening system).

**Figure S2.** Identification of myoDNs. (**A**) Ratio of MHC$^+$ cells within the other lot of mMBs treated with 10 µM PS-ODNs in GM for 48 h (screening system). ** $p < 0.01$ vs control (Dunnett's test). $n = 3$. (**B**) Relative number of the DAPI$^+$ cells of the mMBs shown in panel A. The mean value of the control samples was set at 1.0. ** $p < 0.01$ vs control (Dunnett's test). $n = 3$. (**C**) Ratio of MHC$^+$ cells within the mMBs treated with 10 µM of iSN04' or iSN04 in GM for 48 h (screening system). ** $p < 0.01$ vs control; NS, no significant difference (Scheffe's $F$ test). $n = 3$. (**D**) Representative immunofluorescent images of the C2C12 cells treated with 10 µM iSN04 in DM for 48 h. Scale bar, 200 µm. Ratio of MHC$^+$ cells and multinuclear myotubes were quantified. * $p < 0.05$, ** $p < 0.01$ vs control (Student's $t$ test). $n = 8$. (**E**) Sequences of myoDNs, iSN01-07. Telomeric sequences are underlined. (**F**) Ratio of MHC$^+$ cells within the mMBs treated with 10 µM of iSN02 or denatured-iSN02 (de-iSN02) in GM for 48 h (screening system). PS-ODNs were denatured at 95°C for 5 min then immediately placed on ice. ** $p < 0.01$ vs control, †† $p < 0.01$ vs iSN02 (Scheffe's $F$ test). $n = 3$. (**G**) Ratio of MHC$^+$ cells within the mMBs treated with 10 µM of iSN04 or denatured-iSN04 (de-iSN04) in GM for 48 h (screening system). ** $p < 0.01$ vs control; NS, no significant difference (Scheffe's $F$ test). $n = 3$.

**Figure S3.** iSN04 activity is independent of TLR signal. (**A**) RT-PCR (40 cycles) result of human TLR gene expression in the hMBs maintained in GM. (**B**) RT-PCR (40 cycles) results of murine TLR gene expression in the mMBs and C2C12 cells maintained in GM. (**C**) Heatmap of FPKM values of the genes involved in TLR signaling pathway (KEGG pathway; map04620) in the hMBs treated with 30 μM iSN04 for 24 h. There was no significant difference between control and the iSN04-treated groups in each gene expression (FDR).

**Figure S4.** Profile of iSN04-dependent gene expression. (**A**) Qualities of the RNA samples used for RNA-seq. The RIN values measured by Agilent 2100 Bioanalyzer are 10.0 (max score) in all samples. (**B**) FPKM values of myogenic gene transcripts in the hMBs treated with 30 μM iSN04 for 24 h. * $p < 0.05$, ** $p < 0.01$ vs control (FDR). $n = 3$.

**Figure S5.** Berberine analogs enhance iSN04 activity. (**A**) Structural formula of berberine analogs. (**B**) Representative images of agarose gel electrophoresis of iSN04 mixed with berberine in cationic solutions. The gel was scanned as described in Figure 3E. (**C**) Representative images of agarose gel electrophoresis of iSN01-iSN07 mixed with berberine in F10 medium. (**D**)

Representative images of agarose gel electrophoresis of iSN04 and AS1411 mixed with berberine in F10 medium.

**Figure S6.** iSN04 targets nucleolin and improves p53 protein level. **(A)** Representative immunofluorescent images of the C2C12 cells maintained in DM at day 0 and 2. Scale bar, 50 µm. **(B)** Representative immunofluorescent images of the C2C12 cells treated with 3 µM of iSN04 or AS1411 in DM for 48 h. Scale bar, 50 µm. **(C)** qPCR results of nucleolin gene expression in the hMBs maintained in DM. Mean value of hMBs at day 0 was set to 1.0. ** $p < 0.01$ vs day 0 (Scheffe's $F$ test). $n = 3$. **(D)** FPKM values of nucleolin transcripts in the hMBs treated with 30 µM iSN04 in DM for 24 h. $n = 3$. **(E)** Heatmap of FPKM values of the iSN04-dependent DEGs enriched in the p53 signaling pathway (KEGG pathway; hsa04115) shown in Figure 4E. **(F)** FPKM values of p53 transcripts in the hMBs treated with 30 µM iSN04 in DM for 24 h. ** $p < 0.01$ vs control (FDR). $n = 3$. **(G)** The structure of AS1411 determined by solution NMR (Protein Data Bank, 2N3M). **(H)** Representative immunofluorescent image of MEFs. Scale bar, 50 µm. **(I)** Western blotting (5 µg protein/lane) of nucleolin and GAPDH in the soluble whole cell lysates of C2C12 and MC3T3-E1 cells used for precipitation assay.

**Supplementary Movie.** Simulated conformations of iSN04 at 310 K. 1737 conformations of iSN04 at 310 K were sequentially arrayed to make the movie. These conformations were taken from production run (see Materials and methods in main text) by metropolis method. In the movie, hydrogen atoms were hidden to see the conformations clearly. The colors indicate the residues from 5' end in blue to 3' end in red as rainbow manner.

**Figure S1**

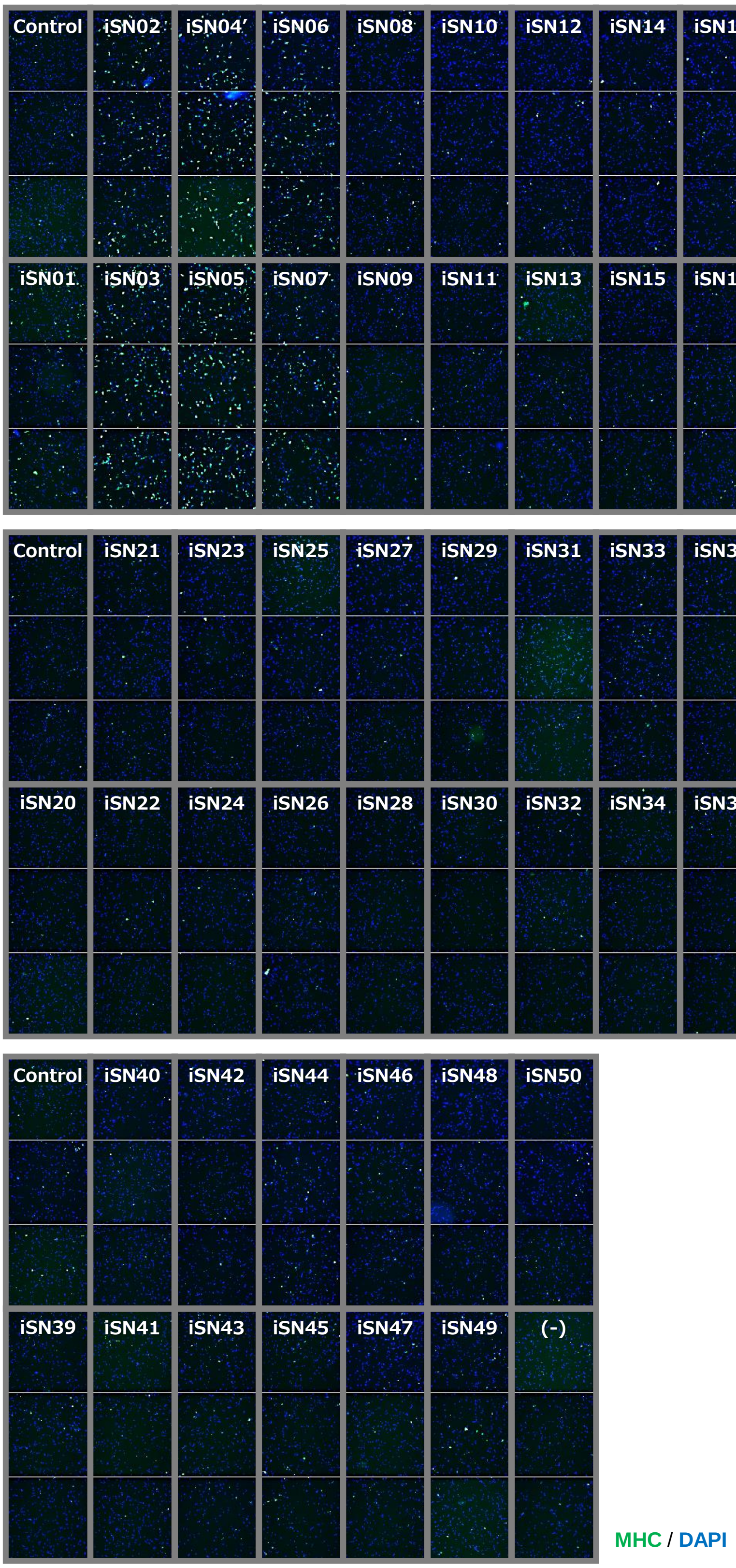
Control
iSN02
iSN04'
iSN06
iSN08
iSN10
iSN12
iSN14
iSN16
iSN18
iSN01
iSN03
iSN05
iSN07
iSN09
iSN11
iSN13
iSN15
iSN17
iSN19
Control
iSN21
iSN23
iSN25
iSN27
iSN29
iSN31
iSN33
iSN35
iSN37
iSN20
iSN22
iSN24
iSN26
iSN28
iSN30
iSN32
iSN34
iSN36
iSN38
Control
iSN40
iSN42
iSN44
iSN46
iSN48
iSN50
iSN39
iSN41
iSN43
iSN45
iSN47
iSN49
(-)
MHC / DAPI

### Figure S2

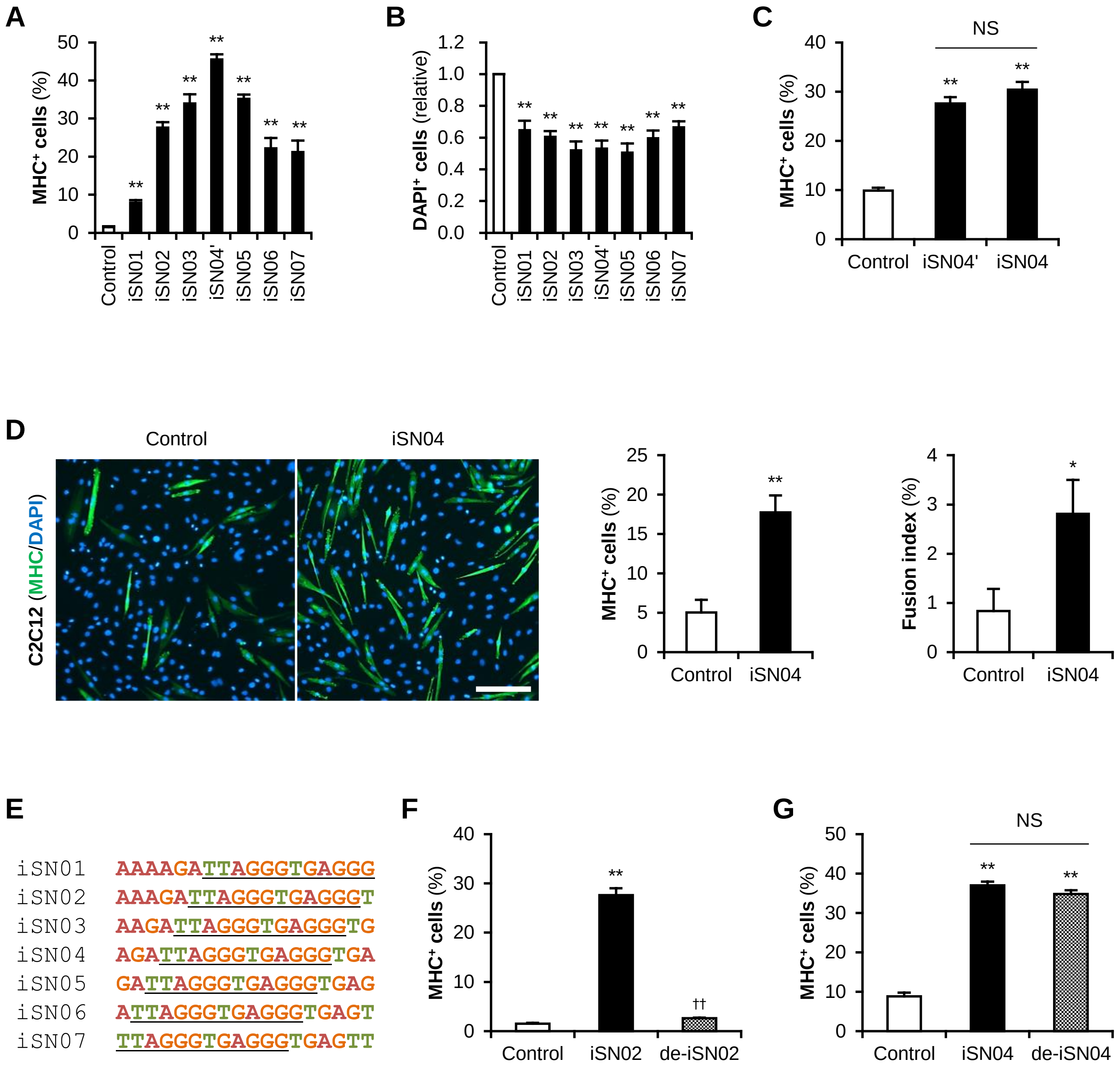
A
MHC+ cells (%)
50
40
30
20
10
0
Control
iSN01
iSN02
iSN03
iSN04'
iSN05
iSN06
iSN07
B
DAPI+ cells (relative)
1.2
1.0
0.8
0.6
0.4
0.2
0.0
C
NS
MHC+ cells (%)
Control
iSN04'
iSN04
D
Control
iSN04
C2C12 (MHC/DAPI)
MHC+ cells (%)
25
20
15
10
5
0
Fusion index (%)
4
3
2
1
0
E
iSN01 AAAAGATTAGGGTGAGGG
iSN02 AAAGATTAGGGTGAGGGT
iSN03 AAGATTAGGGTGAGGGTG
iSN04 AGATTAGGGTGAGGGTGA
iSN05 GATTAGGGTGAGGGTGAG
iSN06 ATTAGGGTGAGGGTGAGT
iSN07 TTAGGGTGAGGGTGAGTT
F
MHC+ cells (%)
iSN02
de-iSN02
G
MHC+ cells (%)
de-iSN04
**
*
††

**Figure S3**

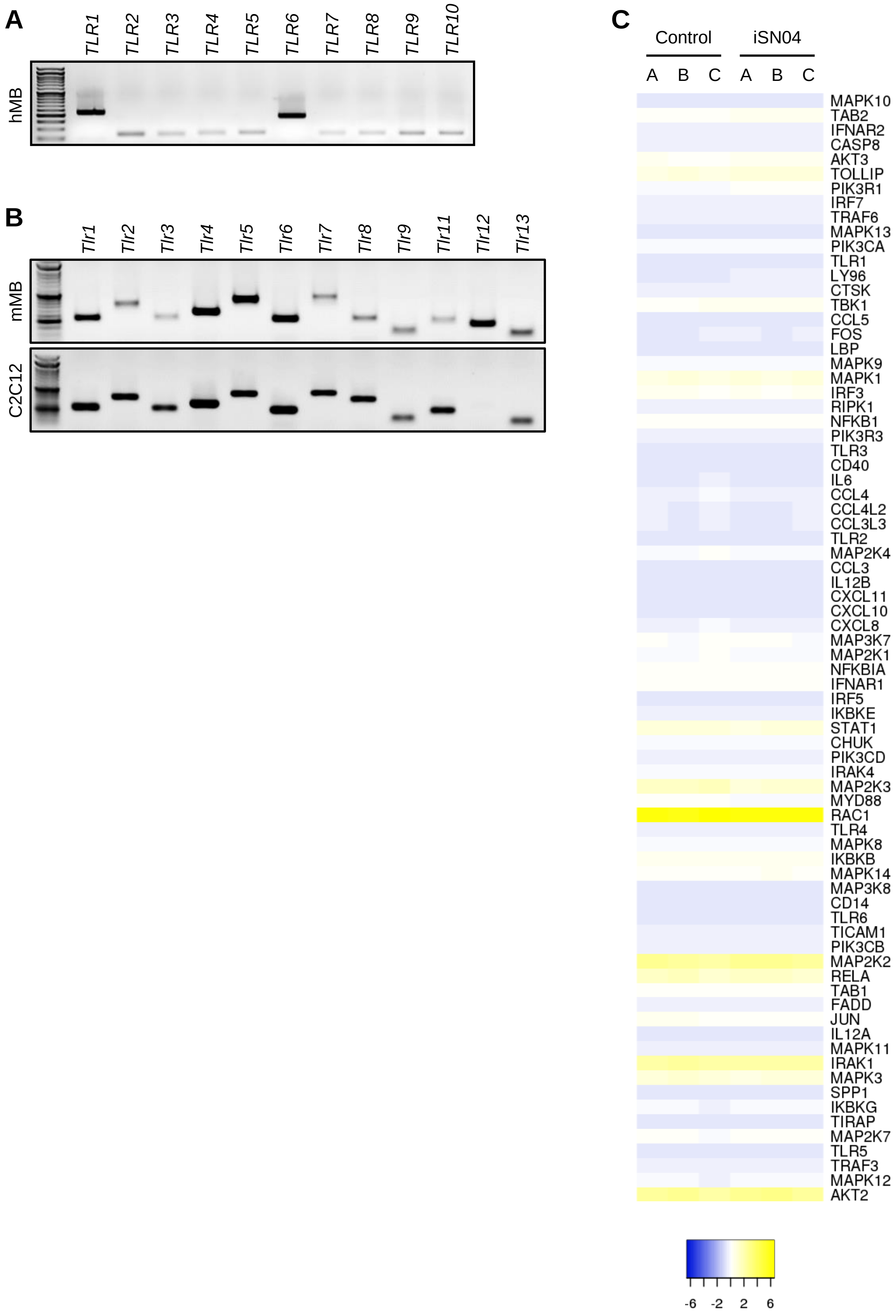
A
TLR1
TLR2
TLR3
TLR4
TLR5
TLR6
TLR7
TLR8
TLR9
TLR10
hMB
B
Tlr1
Tlr2
Tlr3
Tlr4
Tlr5
Tlr6
Tlr7
Tlr8
Tlr9
Tlr11
Tlr12
Tlr13
mMB
C2C12
C
Control
iSN04
A
B
C
A
B
C
MAPK10
TAB2
IFNAR2
CASP8
AKT3
TOLLIP
PIK3R1
IRF7
TRAF6
MAPK13
PIK3CA
TLR1
LY96
CTSK
TBK1
CCL5
FOS
LBP
MAPK9
MAPK1
IRF3
RIPK1
NFKB1
PIK3R3
TLR3
CD40
IL6
CCL4
CCL4L2
CCL3L3
TLR2
MAP2K4
CCL3
IL12B
CXCL11
CXCL10
CXCL8
MAP3K7
MAP2K1
NFKBIA
IFNAR1
IRF5
IKBKE
STAT1
CHUK
PIK3CD
IRAK4
MAP2K3
MYD88
RAC1
TLR4
MAPK8
IKBKB
MAPK14
MAP3K8
CD14
TLR6
TICAM1
PIK3CB
MAP2K2
RELA
TAB1
FADD
JUN
IL12A
MAPK11
IRAK1
MAPK3
SPP1
IKBKG
TIRAP
MAP2K7
TLR5
TRAF3
MAPK12
AKT2
-6
-2
2
6

**Figure S4**

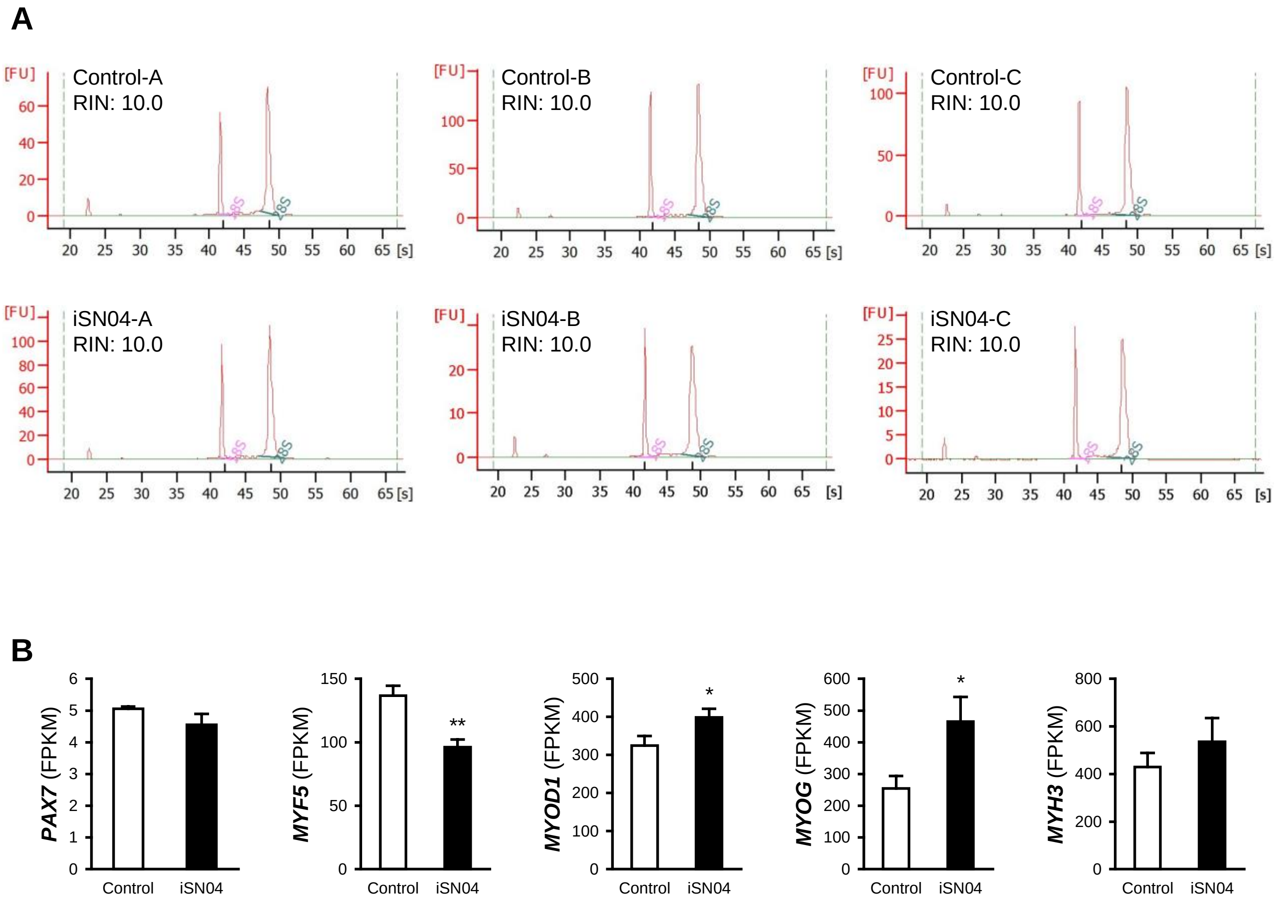
A
Control-A
RIN: 10.0
Control-B
RIN: 10.0
Control-C
RIN: 10.0
iSN04-A
RIN: 10.0
iSN04-B
RIN: 10.0
iSN04-C
RIN: 10.0
[FU]
[s]
18S
28S
B
PAX7 (FPKM)
MYF5 (FPKM)
MYOD1 (FPKM)
MYOG (FPKM)
MYH3 (FPKM)
Control
iSN04
**
*

**Figure S5**

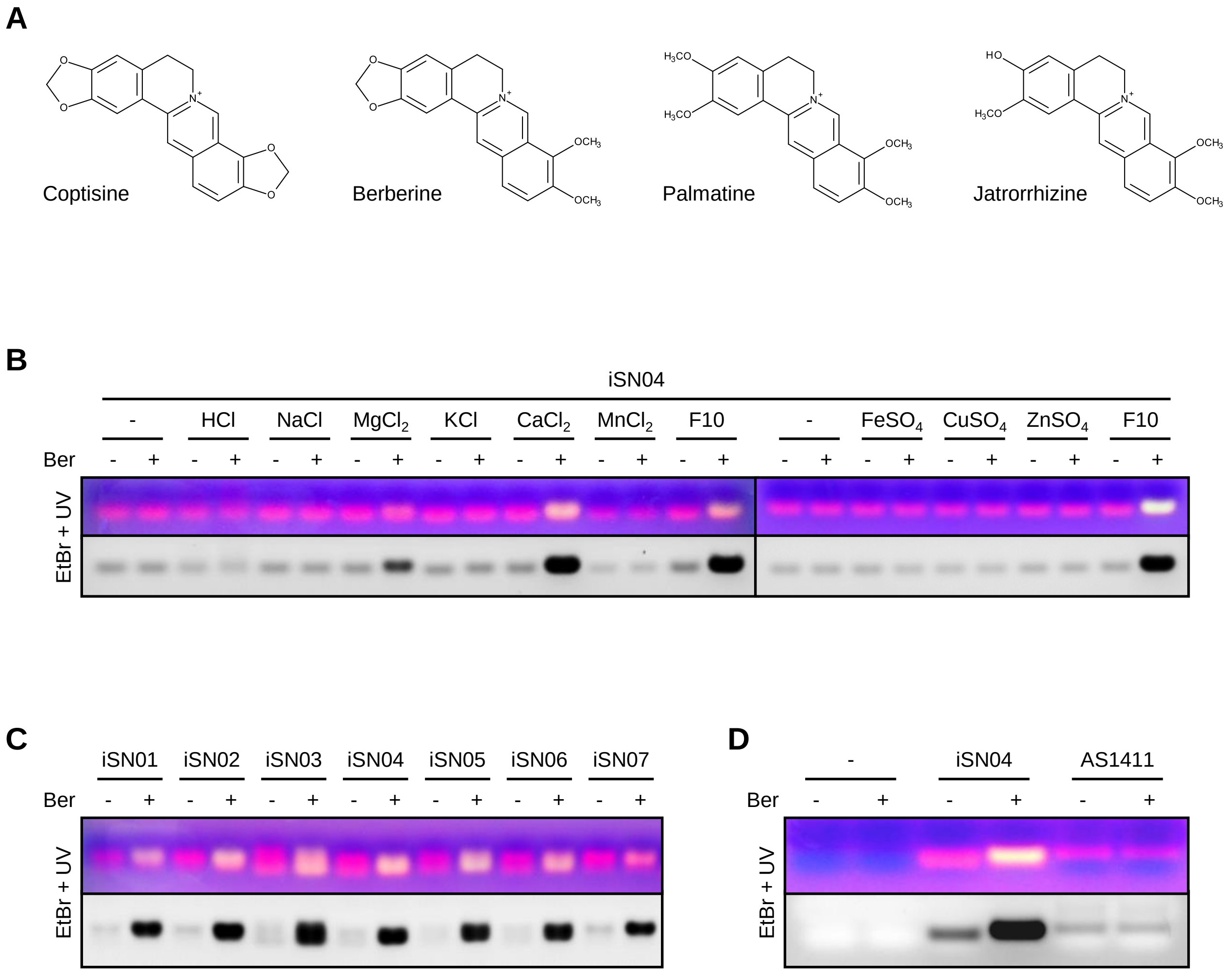
A
Coptisine
Berberine
Palmatine
Jatrorrhizine
O
N+
OCH3
H3CO
HO
B
iSN04
-
HCl
NaCl
MgCl2
KCl
CaCl2
MnCl2
F10
FeSO4
CuSO4
ZnSO4
Ber
EtBr + UV
C
iSN01
iSN02
iSN03
iSN04
iSN05
iSN06
iSN07
D
AS1411
+

### Figure S6

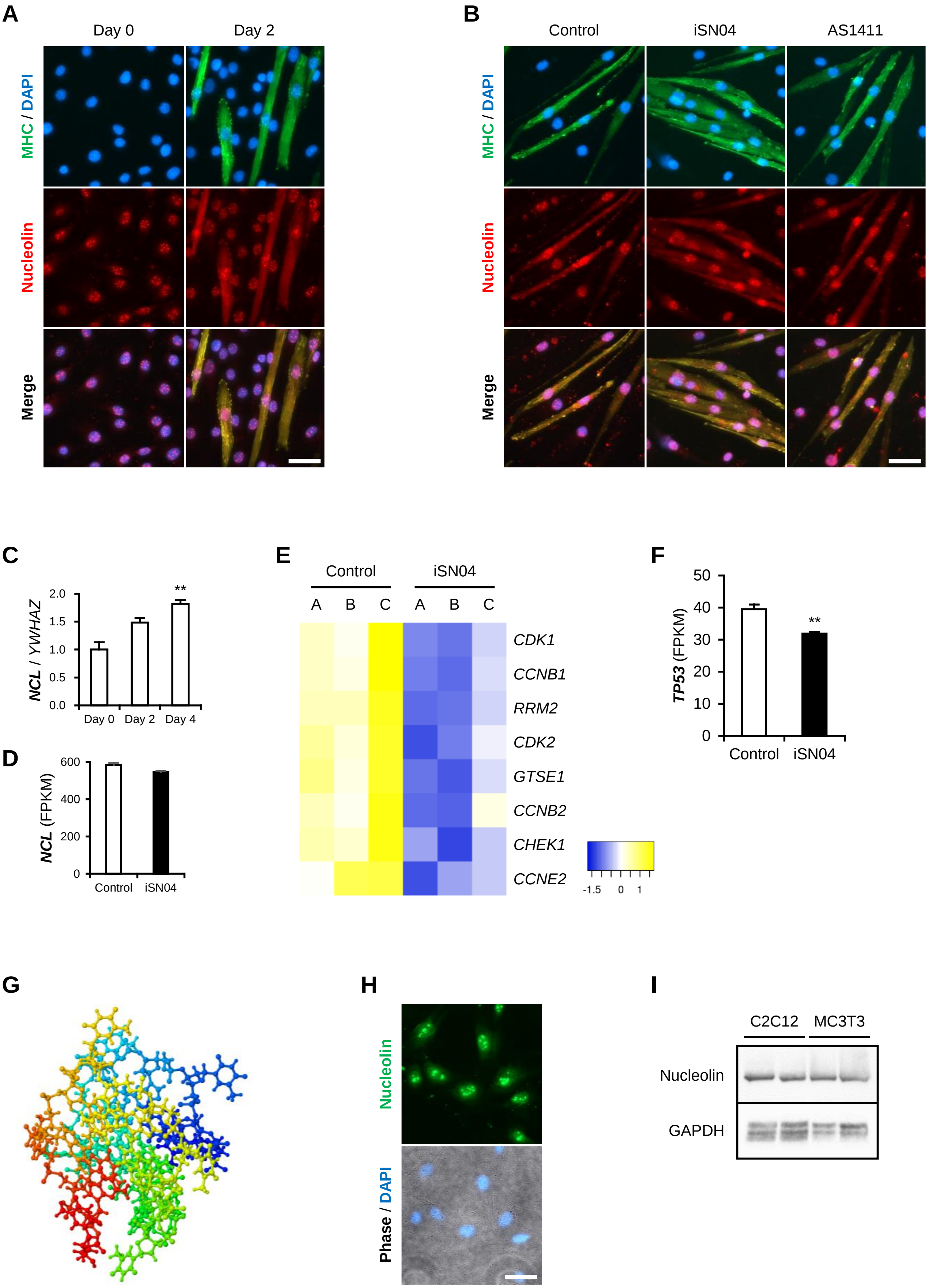

A
Day 0
Day 2
MHC / DAPI
Nucleolin
Merge
B
Control
iSN04
AS1411
MHC / DAPI
Nucleolin
Merge
C
NCL / YWHAZ
2.0
1.5
1.0
0.5
0.0
**
Day 0
Day 2
Day 4
D
NCL (FPKM)
600
400
200
0
Control
iSN04
E
Control
iSN04
A
B
C
A
B
C
CDK1
CCNB1
RRM2
CDK2
GTSE1
CCNB2
CHEK1
CCNE2
-1.5
0
1
F
TP53 (FPKM)
50
40
30
20
10
0
**
Control
iSN04
G
H
Nucleolin
Phase / DAPI
I
C2C12
MC3T3
Nucleolin
GAPDH
